# The role of heparan sulfate in enhancing the chemotherapeutic response in triple-negative breast cancer

**DOI:** 10.1101/2023.09.08.556819

**Authors:** Jasmine M Manouchehri, Lynn Marcho, Mathew A Cherian

**Affiliations:** Comprehensive Cancer Center, The Ohio State University, 410 W 10th Ave, 43210, Columbus, OH, United States

**Author notes:** corresponding author, To whom correspondence should be addressed: Mathew A Cherian, Dept. of Medicine, The Ohio State University Wexner Medical Center, 410 W 10th Ave., Columbus, OH 43210.

## Abstract

**Background:** Among women worldwide, breast cancer has the highest incidence and is the leading cause of cancer-related death. Patients with the triple-negative breast cancer (TNBC) subtype have an inferior prognosis in comparison to other breast cancers because current therapies do not facilitate long-lasting responses. Thus, there is a demand for more innovative therapies that induce durable responses.

In our previous research, we discovered that augmenting the concentration of extracellular ATP (eATP) greatly enhances the chemotherapeutic response of TNBC cell lines by activating purinergic receptors (P2RXs), leading to cell death through the induction of non-selective membrane permeability. However, eATP levels are limited by several classes of extracellular ATPases. One endogenous molecule of interest that can inhibit multiple classes of extracellular ATPases is heparan sulfate. Polysulfated polysaccharide heparan sulfate itself is degraded by heparanase, an enzyme that is known to be highly expressed in various cancers, including breast cancer. Heparan sulfate has previously been shown to regulate several cancer-related processes such as fibroblast growth factor signaling, neoangiogenesis by sequestering vascular endothelial growth factors in the extracellular matrix, hedgehog signaling and cell adhesion. In this project, we identified an additional mechanism for a tumor suppressor role of heparan sulfate: inhibition of extracellular ATPases, leading to augmented levels of eATP.

Several heparanase inhibitors have been previously identified, including OGT 2115, suramin, PI-88, and PG 545. We hypothesized that heparanase inhibitors would augment eATP concentrations in TNBC by increasing heparan sulfate in the tumor microenvironment, resulting in enhanced cell death in response to chemotherapy.

**Methods:** We treated TNBC cell lines MDA-MB 231, Hs 578t, and MDA-MB 468 and non-tumorigenic immortal mammary epithelial MCF-10A cells with increasing concentrations of the chemotherapeutic agent paclitaxel in the presence of heparan sulfate and/or the heparanase inhibitor OGT 2115 while analyzing eATP release and cell viability. Moreover, to verify that the effects of OGT 2115 are mediated through eATP, we applied specific antagonists to the purinergic receptors P2RX4 and P2RX7. In addition, the protein expression of heparanase was compared in the cell lines by Western blot analysis. We also evaluated the consequences of this therapeutic strategy on the breast cancer-initiating cell population in the treated cells using flow cytometry and tumorsphere formation efficiency assays.

**Results:** Heparanase was found to be highly expressed in immortal mammary epithelial cells in comparison to TNBC cell lines. The heparanase inhibitor OGT 2115 augmented chemotherapy-induced TNBC cell death and eATP release.

**Conclusion:** These results demonstrate that inhibiting the degradation of heparan sulfate in the tumor microenvironment augments the susceptibility of TNBC cell lines to chemotherapy by increasing extracellular ATP concentrations. This strategy could potentially be applied to induce more enhanced and enduring responses in TNBC patients.

## Background

Millions of women are affected by breast cancer each year. It was the most common cause of cancer-related mortality among women in 2020 along with the highest global incidence rate with 47.8 new cases and 13.6 deaths per 100,000 per year [1]. Currently, there is a deficit in the availability of specific targeted therapies and a worse outlook for those patients diagnosed with triple-negative breast cancer (TNBC) as compared to other breast cancer subtypes because of the need for progressively more intrusive and toxic therapies to maintain disease control [2–4]. Thus, the development of more efficacious therapies is needed.

There is a noticeable difference in extracellular adenosine triphosphate (eATP) concentrations in cancers [5–7]. Under physiological conditions, the concentration of intracellular ATP can be between 3 to 10 millimolar (mM); whereas the concentration of eATP is between 0-10 nanomolar (nM), a 10^6^-fold difference [8]. Despite this, the minute concentration of eATP can act as a signaling molecule through cell surface purinergic receptors [5–7]. Our previously published study demonstrated that eATP is toxic (in the high micromolar range) to TNBC cells but not to non-tumorigenic immortal mammary epithelial MCF-10A cells [9]. However, eATP can be broken down by different ecto-nucleotidases including ecto-nucleoside triphosphate diphosphohydrolases (E-NTPDase), 5’ nucleotidases, ecto-nucleotide pyrophosphatases/phosphodiesterases (E-NPPase), and tissue non-specific alkaline phosphatases (TNAP). E-NTPDases are considered to be the main enzyme responsible for ATP degradation with extracellular 5’nucleotidase responsible for the catalytic conversion of AMP to adenosine [10]. We previously showed that inhibitors of each of these families of ecto-ATPases possess the capacity to enhance eATP release through the P2RX4 and P2RX7 ion-coupled purinergic receptors. Thus, all ecto-ATPases may need to be inhibited to maximize eATP release and TNBC cell death. The presence of multiple families of ecto-ATPase inhibitors complicates the design of synthetic inhibitors. As part of our interest in broad-spectrum ecto-ATPase inhibitors, we learned that polysulfated polymers such as heparan sulfate inhibit multiple classes of ecto-ATPases [11, 12]. Hence, we hypothesized that enhancement of heparan sulfate levels would exacerbate chemotherapy-induced eATP release and TNBC cell death.

The polysulfated polysaccharide heparan sulfate is synthesized in the Golgi system and is composed of disaccharide units that are negatively charged and unbranched, with sulfation on 3-O, 6-O or N sites of glucosamine as well as the 6-O site on glucuronic/iduronic acid [13–16].

Heparan sulfate impacts growth factor signaling, regulates cell adhesion, and sequesters growth factors in the extracellular matrix (ECM) [14, 15, 17]. Heparan sulfate proteoglycans also modulate the Hedgehog pathway, which is necessary for embryogenesis and tissue homeostasis [18]. Fibroblast growth factors (FGFs) and vascular endothelial growth factors (VEGFs), whose expression levels are elevated in tumors compared to normal tissue, are sequestered by heparan sulfate proteoglycans; FGFs can promote tumor proliferation and VEGFS can regulate angiogenesis [19]. Hence, heparan sulfate may exert its tumor suppressor properties by pathways that are independent of eATP as well.

Heparan sulfate is also involved in inflammatory actions such as retaining cytokines/chemokines in the pericellular microenvironment and promoting the initiation of innate immune responses [15, 17]. Heparanase, the enzyme that degrades heparan sulfate, localizes in the nucleus, lysosomes and late endosomes, and elevated levels of this enzyme have been observed in a variety of cancers, including breast cancer [13–16, 20, 21]. The growth of tumor xenografts is inhibited in the presence of heparanase siRNA, supporting a protumorigenic role for this enzyme [15]. A study that measured levels of heparanase activity in oral cancer cell lines showed that there was a correlation between high heparanase mRNA expression and heparanase activity, suggesting that the expression of the protein is critical for its oncogenic effects [22].

Various heparanase inhibitors have been developed previously, including neutralizing antibodies, peptides and small molecules, such as suramin, PI-88, SST0001, M402 and PG545. These have revealed some effectiveness in vitro and in vivo; PI-88 has been utilized in phase II clinical trials related to prostate cancer while SST0001 has been used in phase I/II clinical trials for multiple myeloma [13–16, 22, 23]. Thus, heparanase inhibition is a potential therapeutic strategy for TNBC.

## Materials and Methods

### Cell culture and drugs and chemicals

Breast cancer cell lines MDA-MB 231 (ATCC HTB-26, RRID: CVCL_0062), MDA-MB 468 (ATCC HTB-132, RRID: CVCL_0419), and Hs 578t (ATCC HTB-126, RRID: CVCL_0332), as well as non-tumorigenic immortalized mammary epithelial MCF-10A cells (ATCC Cat# CRL-10317, RRID: CVCL_0598), were authenticated and maintained as described previously [9].

The following drugs and chemicals were used: ATP (Sigma), paclitaxel (Calbiochem), OGT 2115 (Tocris), A438079 (Tocris), 5-BDBD (Tocris) and heparan sodium sulfate (Sigma).

Heparan sulfate and ATP were dissolved in nuclease-free water (Invitrogen); paclitaxel, OGT 2115, A438079 and 5-BDBD were dissolved in dimethyl sulfoxide/DMSO (Sigma). Table 1 shows drugs’ concentrations and functions; we optimized the drug concentrations that were used for the different assays, starting with the previously used drug concentrations as starting points [24–28]. Drugs were added to media at designated concentrations and applied to cells in an *in vitro* system.

**Table 1.**

| Drug | Concentration(s) | Function | Concentration reference |
| --- | --- | --- | --- |
| Paclitaxel | 50 and 100 μM | Chemotherapeutic agent | [1] |
| OGT 2115 | 20 μM | Heparanase inhibitor | [2] |
| A437809 | 20 μM | P2RX7 inhibitor | [3] |
| 5-BDBD | 20 μM | P2RX4 inhibitor | [4] |
| Heparan sodium sulfate | 50 μM | developmental processes, angiogenesis, and tumor metastasis. | [5] |

### Western blot analysis

Equal numbers of cell types (TNBC MDA-MB 231, Hs 578t, MDA-MB 468 cells and non-tumorigenic immortal mammary epithelial MCF-10A cells) were seeded and cultured for 48 hours to 70%-80% confluency. Cell supernatants were collected. Total cell lysates were prepared, protein quantification performed, proteins denatured, separated, and transferred as previously described [9]. For quantitation of heparanase in cell supernatants, 100 µg heparanase, 10 µl cell supernatants, or adjusted volume of cell supernatants – volume inversely proportionate to the total protein mass in the corresponding cell lysate to normalize to cellular mass, with the supernatant sample with the least corresponding protein mass in lysate set as 10 µL of loaded volume, were loaded onto the gels. Unadjusted (with 10 µl supernatant) blots were stained with Ponceau S (Thermo Fisher Scientific) to reveal the loading amount of proteins. The membranes were blocked with 5% non-fat milk at room temperature for an hour and incubated overnight at 4°C with a primary antibody: heparanase (1:200 dilution; Novus Biologicals, Cat# NBP303846, RRID: AB_2927437) diluted in 5% non-fat milk. The membranes were washed and developed as described previously [9]. GAPDH (Cell Signaling Technology, Cat #3683, RRID: AB_1642205) was used as a loading control. Densitometry was performed on Licor Image Studio (RRID: SCR_015795). The student’s t-test was applied to the applicable assays to ascertain significance. * represents p<0.05 and ** represents p<0.01 relative to protein expression in MCF-10A cells.

### ELISAs

TNBC cancer cells and MCF-10A cells were grown for 48 hours and supernatants were collected. The basal expression levels of heparanase (Abcam, Cat# ab256401) and heparan sulfate (Lifespan Biosciences, Cat# LS-F22183) were assessed in the examined cell lines via ELISA analysis according to the corresponding manufacturer’s directions.

### Flow cytometry analysis of heparan sulfate

Cell surface expression of heparan sulfate in TNBC cell lines, MDA-MB 231, Hs 578t, MDA-MB 468 cells, and non-tumorigenic immortal mammary epithelial MCF-10A cells was assessed by flow cytometry. Cells were detached with accutase (Thermo Fisher Scientific). One million cells were washed in PBS with 0.05% BSA, stained with heparan sulfate (Bioss Cat# bs-5072R, RRID: AB_10856731) plus goat anti-rabbit IgG (H+L) secondary antibody-FITC (Novus Biologicals, Cat# NB 7168, RRID: AB_524413) or stained with rabbit IgG Isotype Control-FITC (Invitrogen, Cat# PA5-23092, RRID: AB_2540619) in Flow Cytometry Staining Buffer (2% FBS, 0.02% sodium azide and PBS). Analysis was performed on BD FACS Fortessa using the FITC channel (530/30 nm) and Flowjo software (RRID: SCR_008520). The student’s t-test was applied to ascertain significance. * represents p<0.05 and ** represents p<0.01 relative to MFI in MCF-10A cells; + represents p<0.05 and ++ represents p<0.01 relative to MFI of HEK293-empty vector transfected cells. O/E represents overexpressed.

### Immunohistochemistry of heparanase and heparan sulfate

AMSBIO BR1202B breast cancer tissue array (120 core array with 82 TNBC cores) on Fisher Superfrost Plus slides, was sectioned at 5 µm and air dried overnight. Staining was performed at Histowiz, Inc. Brooklyn, NY using the Leica Bond RX automated stainer (Leica Microsystems). Three normal and two ductal carcinoma in situ (DCIS) slides were used as controls. Samples were processed, embedded in paraffin, and sectioned at 4 μm. These control slides were dewaxed using xylene and serial dilutions of ethanol. Epitope retrieval was performed by heat-induced epitope retrieval (HIER) of the formalin-fixed, paraffin-embedded tissue using citrate-based pH 6 solution (Leica Microsystems, Cat#AR9961) for 10 mins at 95°C. The tissues were first incubated with peroxide block buffer (Leica Microsystems, Cat# RE7101-CE), followed by incubation with the rabbit heparanase antibody (Novus Biologicals, Cat# NBP303846, RRID: AB_2927437) or rat anti-heparan sulfate proteoglycan 2/perlecan antibody [A7L6] (Abcam, Cat# ab2501, RRID: AB_2295402) at 1:50 dilution for 30 mins, followed by DAB rabbit secondary reagents: polymer, DAB refine and hematoxylin (Bond Polymer Refine Detection Kit, Leica Microsystems, Cat# DS9800) according to the manufacturer’s protocol. The slides were dried, coverslipped (TissueTek-Prisma Coverslipper) and visualized using a Leica Aperio AT2 slide scanner (Leica Microsystems) at 40X. For the analysis of the tissue microarray (TMA), the Halo TMA module was used to identify and extract the individual TMA cores by means of constructing a grid over the TMA. All subsequent analysis steps were the same for the three slides of DCIS tissue, the two slides of normal breast tissue and the TMA slide for each antibody. In the first part of the analysis, the tumor area was identified by training a random forest classifier algorithm to separate viable tumor tissue from any surrounding stroma and necrosis areas. Once the tumor area was identified the analysis then proceeded to identify positive and negative cells based on heparan sulfate or heparanase staining within the defined tumor area on each slide and each core from the TMA slide. Positive and negative cells were identified using the Halo Multiplex IHC algorithm v3.4.1 by first defining the settings for the hematoxylin counterstain, followed by setting thresholds to detect the heparan sulfate or heparanase stain positivity of weak, moderate and strong intensities (Halo threshold settings 0.11, 0.35, 0.45). H-score was then generated following the convention below: Weak positive (1+), moderate positive (2+), and strong positive (3+). For the histology statistical analysis, group differences were determined using Kruskal-Wallis one-way analysis of variance. When shown to be statistically significant, a post hoc Dunn’s test was done to determine p values. P values were adjusted to account for multiple comparisons and an alpha level of 0.05 was used for all the tests. The software GraphPad Prism version 10.0.2 (RRID: SCR_002798) was used for all tests.

### Verification of heparanase inhibition by OGT 2115 using levels of heparan sulfation of Syndecan-1

MDA-MB 468 cells were seeded and cultured and cell lysates were prepared as described above. OGT 2115 at 20 µM and 40 µM were applied to 100 µg cell lysates in the absence and presence of 50 units of recombinant heparanase enzyme. The negative control was untreated MDA-MB 468 cell lysates (100 µg) and the positive control cell lysates (100 µg) treated with 100 units of heparin, a known heparanase inhibitor. These treated and untreated lysates were all incubated for 6 hours at 37° C. Protein samples (100 µg) were loaded onto the 8% Tris-Glycine gels, transferred, and blocked in 5% non-fat milk for an hour. The membranes were incubated overnight at 4°C with a syndecan-1 primary antibody:(Abcam, Cat#ab128936, RRID: AB_11150990) diluted in 5% non-fat milk. Densitometry using GAPDH as loading control was carried out as described above. The student’s t-test was applied to the applicable assays to ascertain significance. * represents p<0.05 and ** represents p<0.01 relative to expression in control MDA-MB 468 cell lysates.

### Cell viability and eATP assays

TNBC MDA-MB 231, Hs 578t, MDA-MB 468 cells and non-tumorigenic immortal mammary epithelial MCF-10A cells were plated as previously described and treated with paclitaxel (vehicle), heparan sodium sulfate (50 µM), OGT 2115 (20 µM), A438709 (20 µM), 5-BDBD (20 µM), or different combinations of these drugs. Cells were treated with OGT 2115 and heparan sodium sulfate for 48 hours and with paclitaxel, A438709 or 5-BDBD for the final 6 hours of the 48-hour time course (we treated cells with paclitaxel for 6 hours to replicate exposure times in patients), and cell viability was assessed by applying the PrestoBlue™ HS cell viability reagent (Invitrogen) in accordance with the manufacturer’s instructions [9]. ATP was assessed in supernatants as described above. Fluorescence readings (excitation and emission ranges: 540– 570 nm and 580–610 nm) were assessed using a Bioteck Synergy HT plate reader. One-way ANOVA with Tukey’s HSD (Honestly Significant Difference) was calculated to ascertain significance. * represents p<0.05 and ** represents p<0.01 when comparing vehicle addition to OGT 2115, to heparan sulfate, and the combination of heparan sulfate, OGT 2115 and/or paclitaxel.

### Cancer-initiating cell formation

Breast cancer-initiating cells express aldehyde dehydrogenase (ALDH) and CD44 but not CD24 [29–32]. TNBC MDA-MB 231, Hs 578t and MDA-MB 468 cells were collected and stained following the protocol for the Aldeflour Kit (STEMCELL, Cat#01700). Cells were washed and stained with CD24-PE (eBioscience, Cat# 12-0247-42, RRID: AB_1548678), CD44-APC (eBioscience, Cat# 17-0441-82, RRID: AB_469390) and LIVE/DEAD™ Fixable Near-IR Dead Cell Stain Kit (Invitrogen, Cat# L10119) for 30 minutes at 4°C. Cells were washed and resuspended in the Aldefluor Buffer provided in the Aldeflour kit. Flow cytometry analysis was performed on BD FACS Fortessa using the FITC (ALDH), PE (CD24), APC (CD44) and AF750 (Live/Dead) channels and applying the Flowjo software (RRID: SCR_008520). For statistical analysis, * represents p<0.05 and ** represents p<0.01 when comparing paclitaxel to paclitaxel and OGT 2115.

### Tumorsphere formation efficiency assay

TNBC MDA-MB 231, Hs 578t and MDA-MB 468 cell lines were grown, treated with paclitaxel, OGT 2115 and/or heparan sodium sulfate as described above in the cell viability and eATP section. Cells were trypsinized, washed, resuspended in 3D Tumorsphere Medium XF (Sigma), and plated at 10 viable cells per well after (45 µM) filtration. Cells were grown for seven days, and tumorspheres were counted for each different condition using the Etaluma™ Lumascope 620. One-way ANOVA with Tukey’s HSD was calculated to ascertain significance. For significance, ** represents p<0.01 when comparing paclitaxel to paclitaxel and OGT 2115.

## Results

### Analysis of heparanase and heparan sulfate expressions in breast cancer

#### Heparan sulfate expression by ELISAs, immunochemistry and flow cytometry in breast cancer cell lines and immortal mammary epithelial cells

TNBC MDA-MB 231, Hs 578t and MDA-MB 468 cell lines and non-tumorigenic immortal epithelial mammary MCF-10A cells were examined for extracellular expression of heparan sulfate via ELISAs (Figure 1A) and immunohistochemistry (Figure 1C and 2) and then cell surface expression via flow analysis (Figure 1B). The immortal MCF-10A cell line expressed significantly more heparan sulfate on the cell surface in comparison to the cell surface expression of heparan sulfate amongst the examined TNBC cell lines. These results matched the immunohistochemistry results in that the tissue sections of normal breast and DCIS had significantly higher percentages of cells positive for heparan sulfate than those of invasive breast cancers; however, as stated previously our ELISA results for extracellular heparan sulfate differed as the TNBC MDA-MB 468 cell line expressed significantly more extracellular heparan sulfate than MCF-10A cell line while MCF-10A cells significantly expressed more heparan sulfate than MDA-MB 231 and Hs 578t.

**Figure 1:**
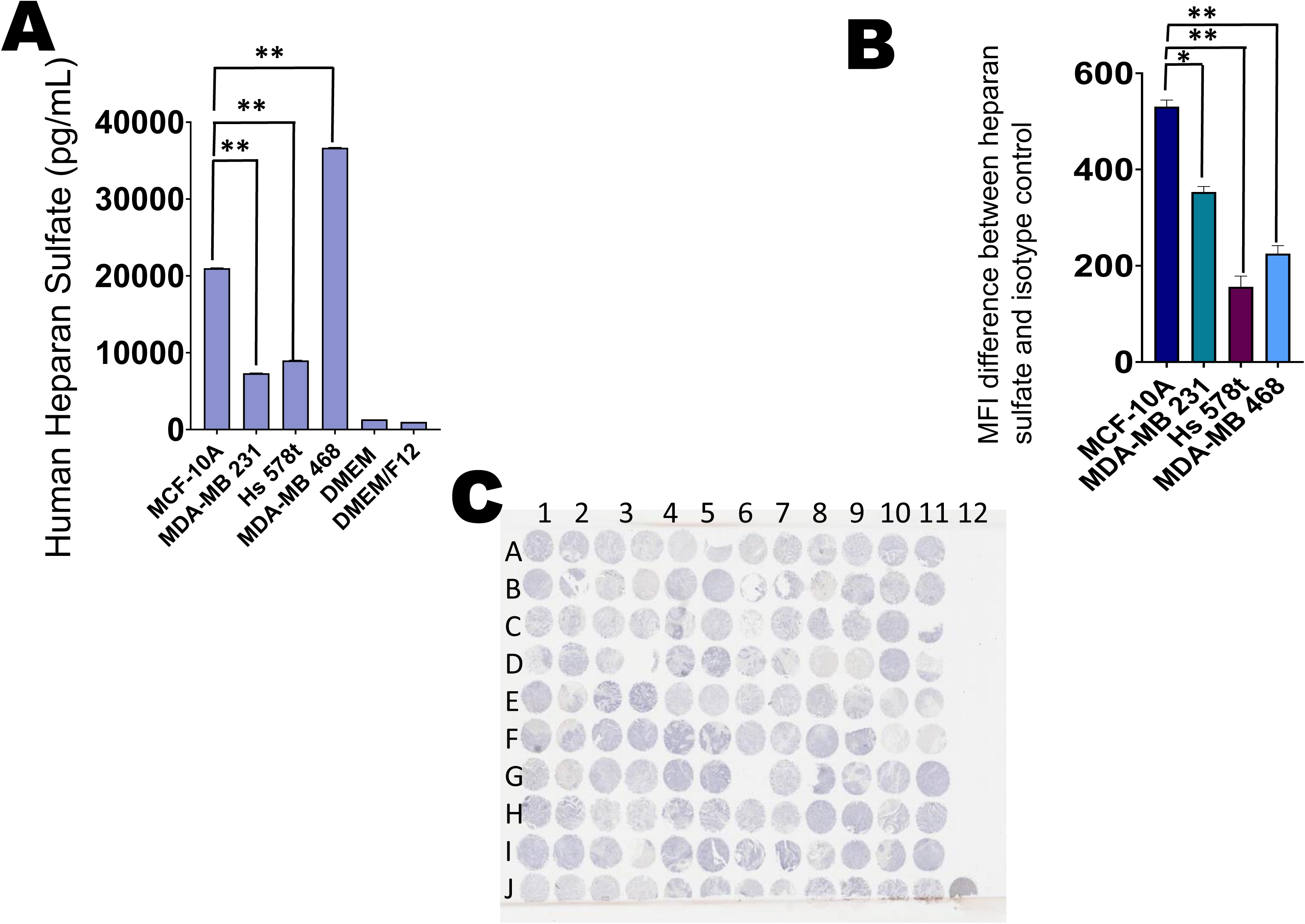
Extracellular and cell surface expression of heparan sulfate. **(A)** Basal heparan sulfate expression was examined in the supernatants of TNBC cells and control immortal MCF-10A via ELISA analysis with MDA-MB 468 expressing the most. The standard deviation was calculated from three independent experiments performed in triplicate. The student’s t-test was performed to determine the significance with * representing p<0.05 and ** representing p<0.01 the protein expression in MCF-10A to the protein expressions in the TNBC cell lines. **(B)** Baseline heparan sulfate cell surface expression levels were examined via flow cytometry analysis in nontumorigenic immortal mammary epithelial MCF-10A cells and TNBC MDA-MB 231, Hs 578t and MDA-MB 468 cells. The standard deviation was calculated from three independent experiments performed in triplicate. The student’s t-test was performed with * representing p<0.05 and ** representing p<0.01 comparing expression levels in MCF-10A to those in the TNBC cell lines. **(C)** The AMSBIO BR1202B breast cancer tissue array (120 cores with 82 TNBC cores; key can be found in Supplemental Figure 8C and corresponding Supplemental Table 1 showing breast cancer sub-type distribution) was stained with heparan sulfate.

**Figure 2:**
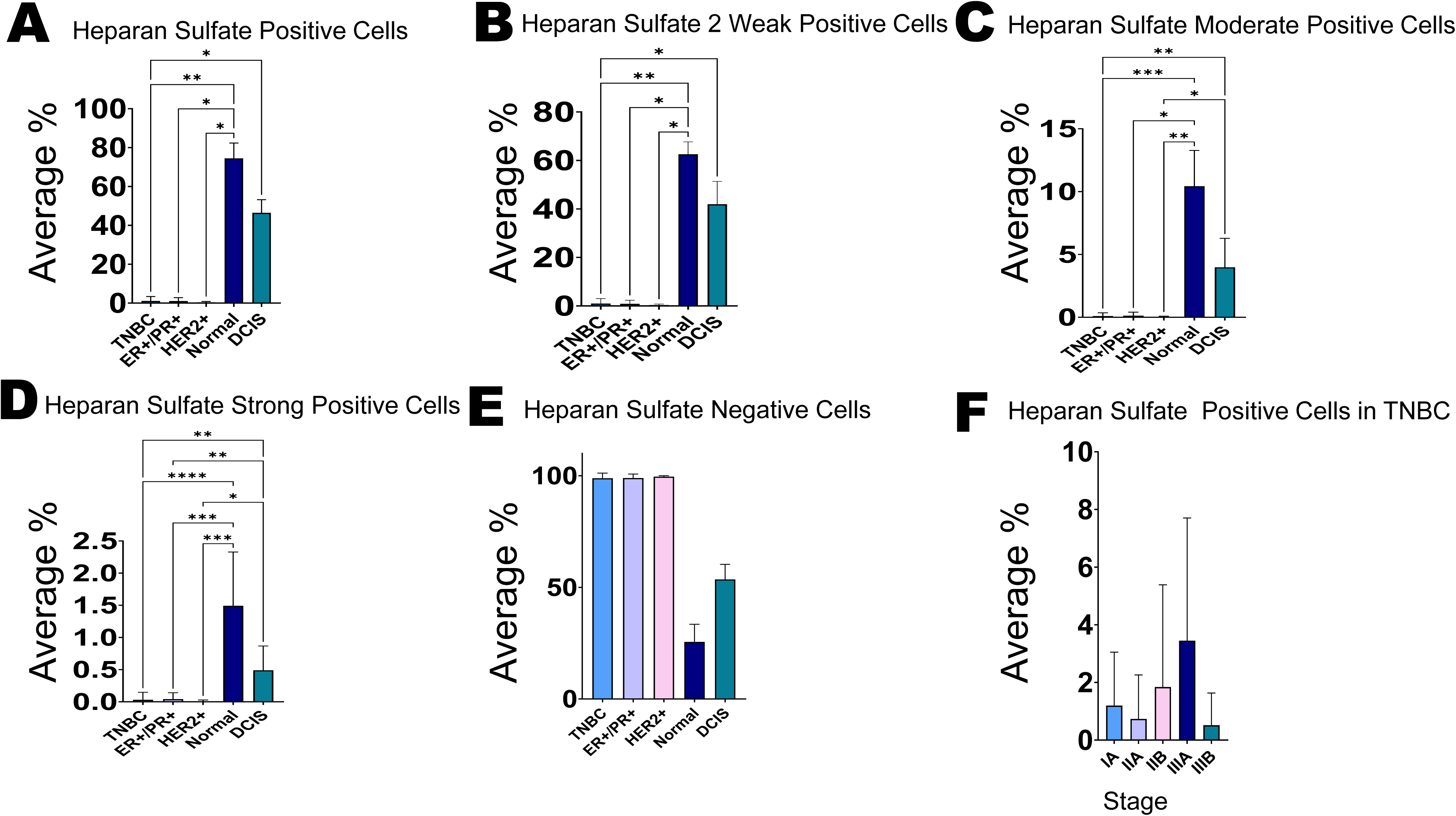
Statistical analysis for heparan sulfate immunohistochemistry comparing different breast cancer subtypes and normal breast tissue. **(A)** Pairwise comparisons using Dunn’s test indicated that there was a significant difference between TNBC and normal breast (p = 0.0040), TNBC and DCIS (p = 0.0359), ER+/PR+ and normal (p = 0.0209), and HER2+ and normal (p = 0.0403), in the percentage of heparan sulfate positively stained cells in the tissue sections. No other differences were significantly different. **(B)** Pairwise comparisons using Dunn’s test showed that there was a significant difference between TNBC and normal (p = 0.0048), TNBC and DCIS (p = 0.0416), ER+/PR+ and normal (p = 0.0131), and HER2+ and normal (p = 0.0321), in the percentage of heparan sulfate weakly stained cells. No other differences were significantly different. **(C)** Pairwise comparisons using Dunn’s test demonstrated that there was a significant difference between TNBC and normal (p = 0.0003), TNBC and DCIS (p = 0.0064), ER+/PR+ and normal (p = 0.0127), HER2+ and normal (p = 0.0041), and HER2+ and DCIS (p = 0.0305), in the percentage of heparan sulfate moderately stained cells. No other differences were significantly different. **(D)** Pairwise comparisons using Dunn’s test demonstrated that there was a significant difference between TNBC and Normal (p = 0.0001), TNBC and DCIS (p = 0.0020), ER+/PR+ and normal (p = 0.0002), ER+/PR+ and DCIS (p = 0.0087), HER2+ and normal (p = 0.0003), and HER2+ and DCIS (p = 0.0109), in the percentage of heparan sulfate strongly stained cells. No other differences were significantly different. **(E)** Pairwise comparisons using Dunn’s test demonstrated that there was a significant difference between TNBC and normal (p = 0.0040), TNBC and DCIS (p = 0.0359), ER+/PR+ and normal (p = 0.0209), HER2+ and normal (p = 0.0003), and HER2+ and normal (p = 0.0403), in the percentage of cells negative for heparan sulfate staining. No other differences were significantly different. **(F)** Kruskal-Wallis test indicated that there was no significant difference in the percentage of heparan sulfate positively stained cells in the tumor amongst the TNBC breast cancer stages.

#### Measurement of heparan sulfate expression in human breast cancer samples by immunohistochemistry

An AMSBIO breast cancer tissue array (120 cores specifically with 82 TNBC cores), two normal breast tissue slides and three DCIS tissue slides were stained with heparan sulfate specific antibody (Figure 1C, Supplemental Figure 8) and statistical analysis was performed (Figure 2). The expression of heparan sulfate was compared amongst TNBCs, estrogen receptor-positive/progesterone positive (ER+/PR+), human epidermal growth factor receptor 2 positive (HER2+) breast cancer, normal breast tissue and DCIS. We carried out a pairwise comparison using Dunn’s test: We found there was a significantly lower percentage of cells that stained positively (any level of staining), weakly, moderately and strongly positive for heparan sulfate in tissue sections of invasive cancers compared to those of normal breast tissue and DCIS. We also carried out the Kruskal-Wallis test which indicated that there was no significant difference between the percentages of cells that stained positively for heparan sulfate amongst TNBC breast cancer stages. Additional statistical analysis was carried out comparing the heparan sulfate expression amongst normal breast tissue, DCIS, and invasive breast cancers. We found that there was a significant difference amongst the different grades of TNBCs and different heparan sulfate stain strengths; normal breast cells expressed the most heparan sulfate (Supplemental Figure 10). Furthermore, we observed no significant difference or pattern when comparing heparan sulfate expression in invasive cancers with different levels of % Ki67 expression (Supplemental Figure 12).

#### Heparanase expression in breast cancer cell lines and mammary epithelial cells by western blot and ELISAs

We wanted to reveal the basal level of expression of heparanase amongst all the examined cell lines. Western blot analysis was performed on TNBC MDA-MB 231, Hs 578t and MDA-MB 468 cell lines and non-tumorigenic immortal epithelial mammary MCF-10A cells, probing for heparanase with GAPDH as the internal loading control (Figure 3). Unexpectedly, TNBC cell lines expressed less heparanase extracellularly when compared to MCF-10A cells as assessed by semi-quantitative densitometry. However, intracellularly heparanase expression was more mixed with MDA-MB 231 expressing the least and Hs 578t and MDA-MB 468 expressing a little less heparanase than MCF-10A. Also, we carried out ELISAs for heparanase expression and saw no difference amongst the examined cell lines using the media for cell lines as a negative control (DMEM for the TNBC cells and DMEM/F12 for MCF-10A) (Figure 4A) which matched the immunohistochemistry results in that there was no significant difference of heparanase expression amongst different invasive breast cancer sub-types and normal breast tissue and DCIS (Figure 4B and Figure 5). However, absolute levels of heparanase must take into account concurrent heparan sulfate expression levels as heparanase has been found to bind to heparan sulfate; thus, heparanase levels positively correlate with and are dependent on heparan sulfate levels [33]. Hence, a more accurate way to analyze heparanase expression may be the ratio of heparanase to heparan sulfate. Thus, when heparan sulfate levels are taken into account (low in TNBC cell lines compared to MCF10A), the paradoxically lower expression of heparanase in TNBC cell lines compared to MCF10A cells may be explained.

**Figure 3:**
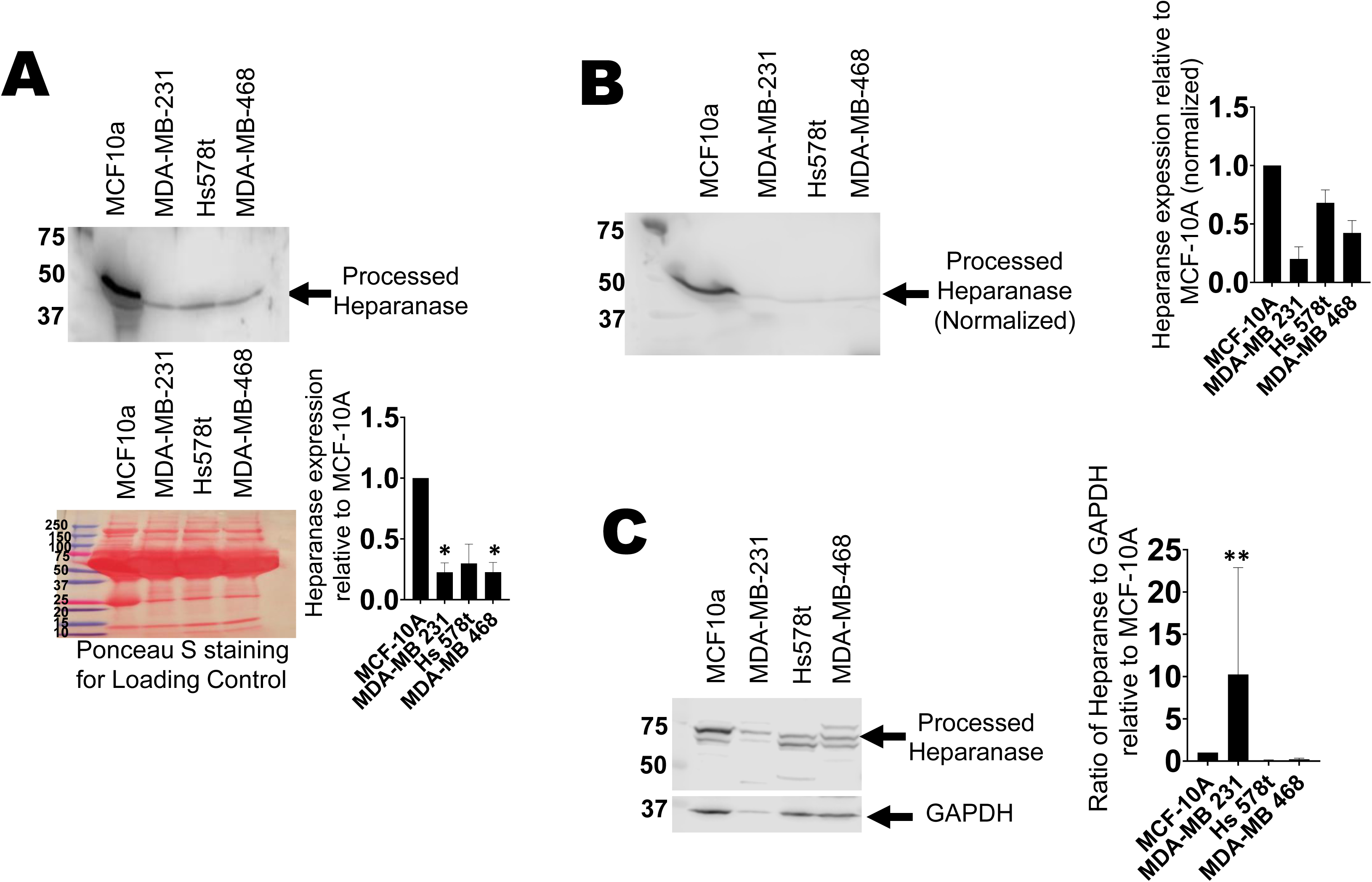
Immunoblot analysis of heparanase expression in immortal mammary epithelial cells and triple-negative breast cancer cell lines. For the western blot analysis of heparanase, **(A)** 10 μl of cell supernatants of nontumorigenic immortal mammary epithelial MCF-10A cells and TNBC MDA-MB 231, Hs 578t and MDA-MB 468 cells were probed with heparanase antibody. **(B)** Adjusted volume of cell supernatants volume inversely proportionate to to the total protein mass in the corresponding cell lysate were probed with heparanase antibody. **(C)** Equal amounts of cell lysate from each cell line were probed. All experiments were repeated once with biological replicates. The densitometric analyses of the bands were calculated. The student’s t-test was performed to determine significance. * represents p<0.05 and ** represents p<0.01 when comparing expression in MCF-10A cells to the expression in TNBC cell lines.

**Figure 4:**
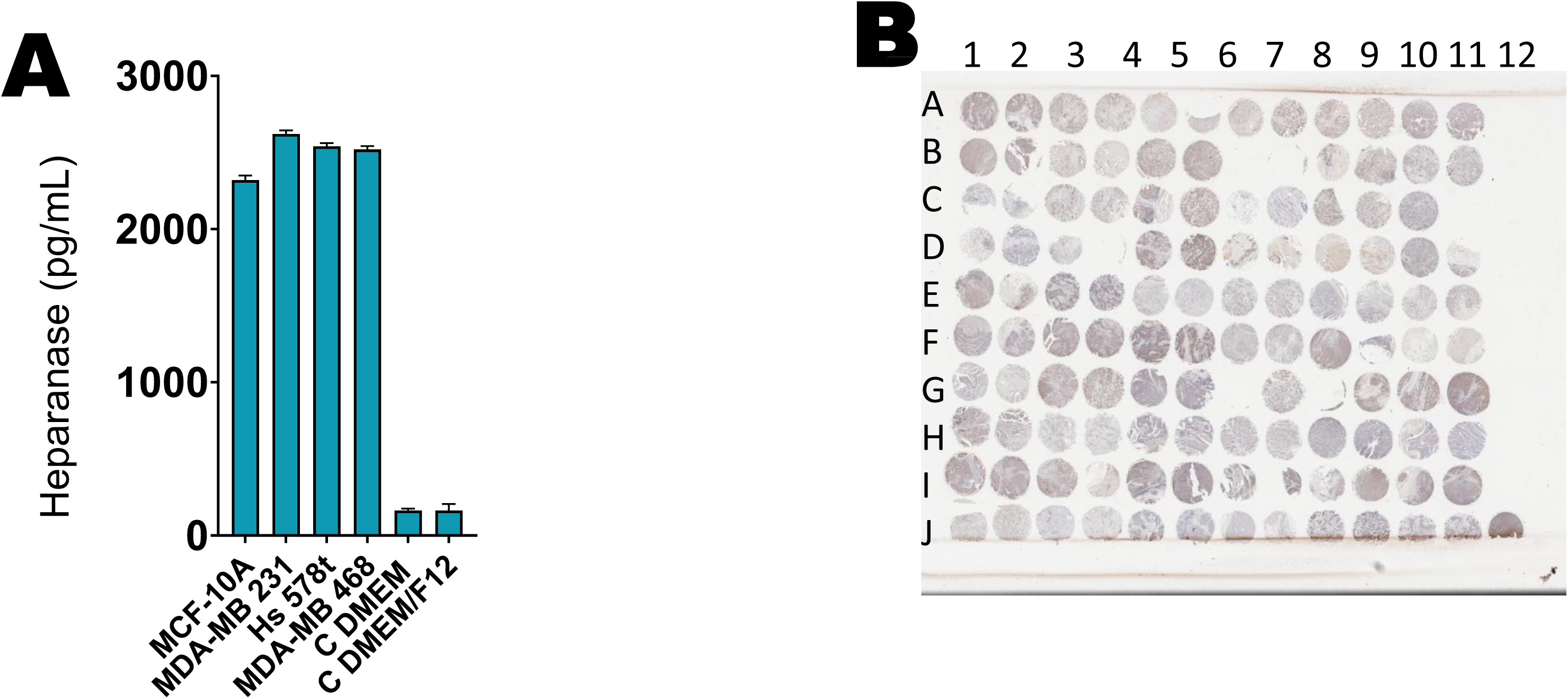
Extracellular expression of heparanase and cell surface expression of heparan sulfate expressions. **(A)** Baseline extracellular heparanase expression was determined in nontumorigenic immortal mammary epithelial MCF-10A cells and TNBC MDA-MB 231, Hs 578t and MDA-MB 468 cells by ELISAs. The standard deviation was calculated from three independent experiments performed in triplicate. The student’s t-test was performed with * representing p<0.05 and ** representing p<0.01, comparing expression levels in MCF-10A to those in the TNBC cell lines. **(B)** The AMSBIO BR1202B breast cancer tissue array (120 cores with 82 TNBC cores; key can be found in Supplemental Figure 8C and corresponding Excel worksheet showing breast cancer sub-type distribution) was stained with heparanase.

**Figure 5:**
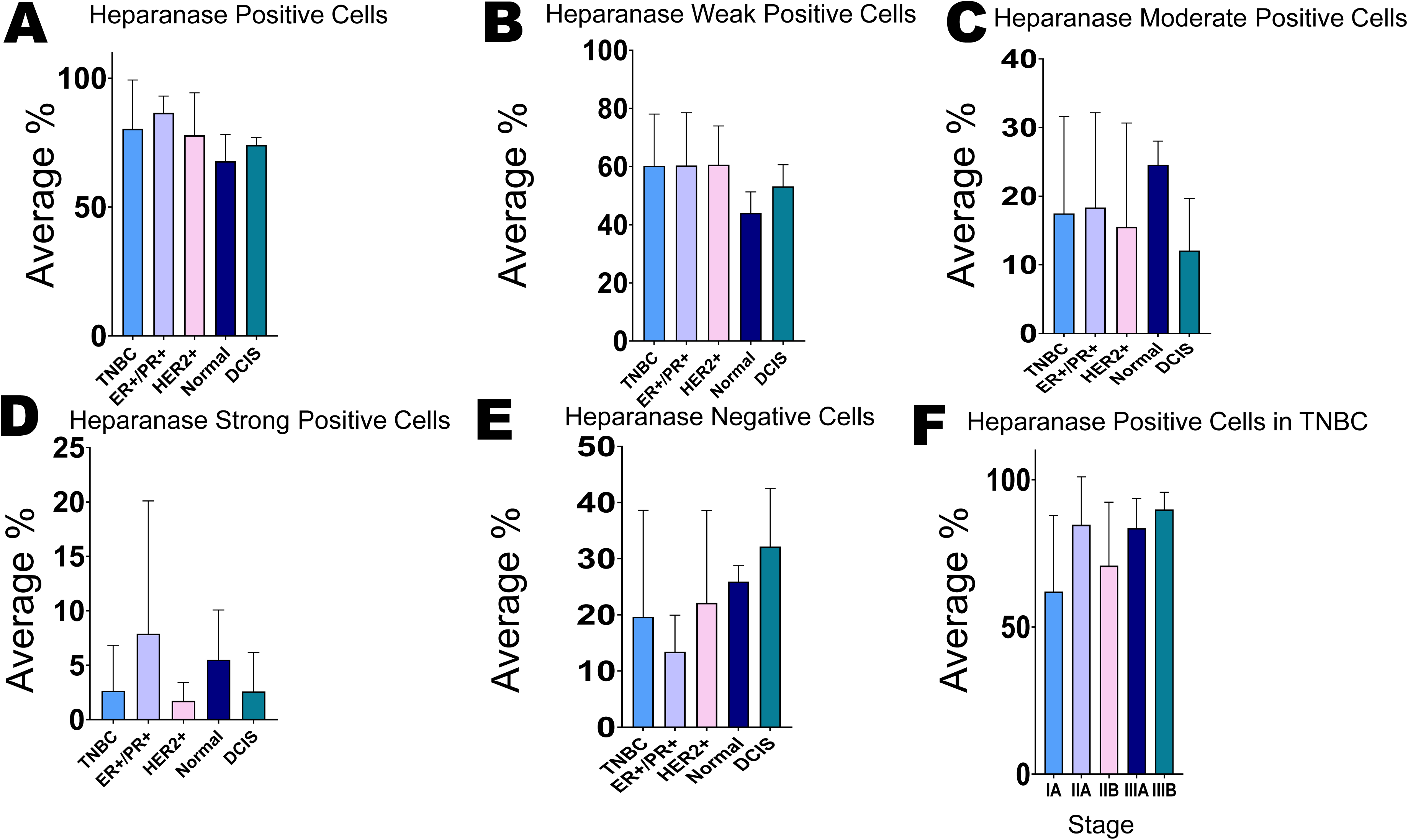
Statistical analysis for heparanase immunohistochemistry comparing different breast cancer subtypes and normal breast tissue. Kruskal-Wallis test indicated that there was no significant difference: **(A)** in the percentage of heparanase positively stained cells in the tissue sections of normal breast tissue, DCIS and invasive breast cancer subtypes, **(B)** in the percentage of heparanase weakly stained cells in the tissue sections of normal breast tissue, DCIS and invasive breast cancer subtypes, **(C)** in the percentage of heparanase moderately stained cells in the tissue sections of normal breast tissue, DCIS and invasive breast cancer subtypes, **(D)** in the percentage of heparanase strongly stained cells in the tissue sections of normal breast tissue, DCIS and invasive breast cancer subtypes, **(E)** in the percentage of negative heparanase stained cells in the tissue sections of normal breast tissue, DCIS and invasive breast cancer subtypes and **(F)** in the percentage of heparanase positively stained cells in the tissue sections of different TNBC breast cancer stages.

We also probed TNBC MDA-MB 231, Hs 578t and MDA-MB 468 cell lines and non-tumorigenic immortal epithelial mammary MCF-10A cells treated with paclitaxel (100 µM) or ATP (500 µM) for heparanase and saw no significant change in expression for heparanase with chemotherapy treatment (Supplemental Figures 4-6).

**Figure 6:**
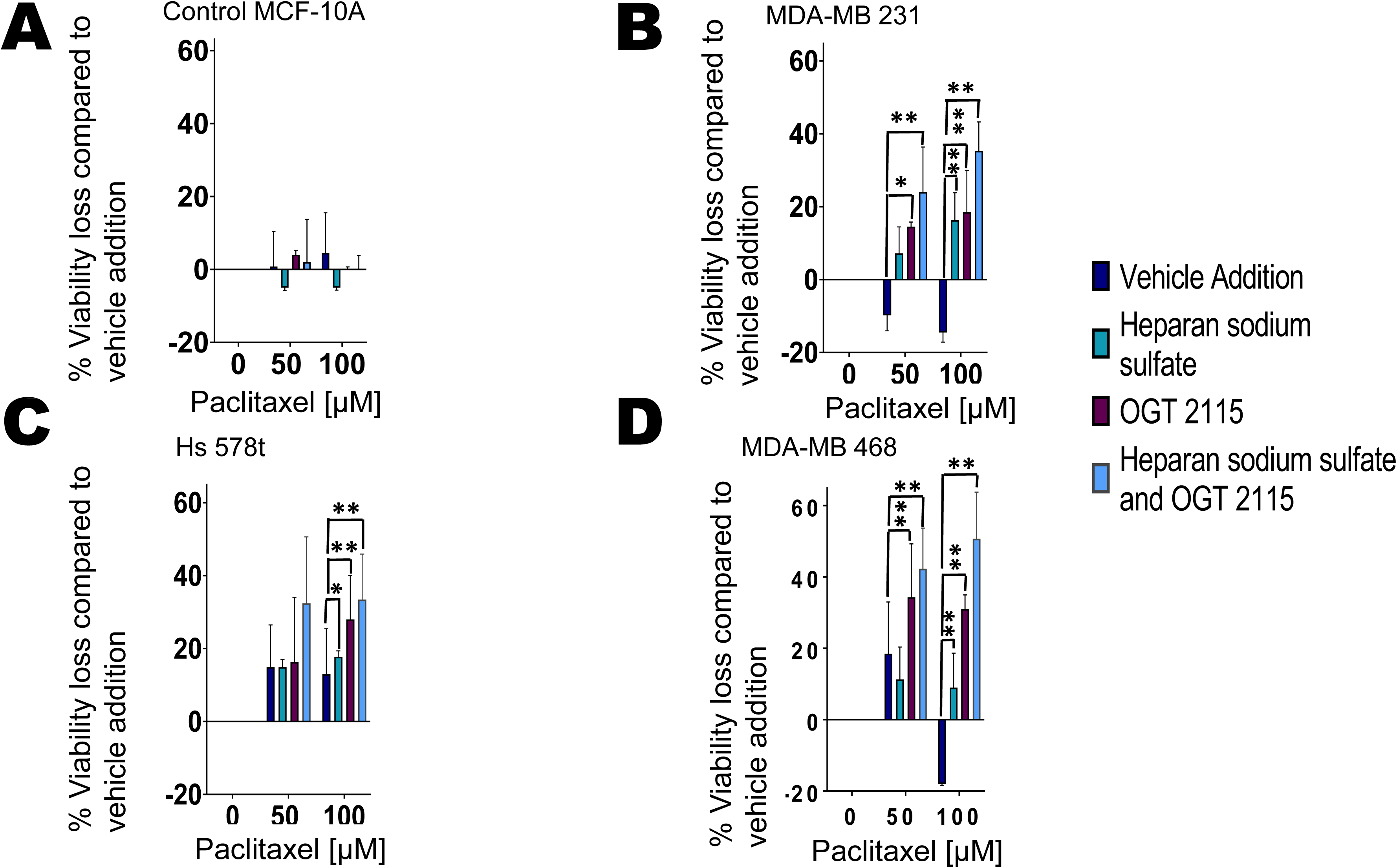
Effects of heparanase inhibitor OGT 2115 and chemotherapeutic agent paclitaxel on cell viability. Percentage loss of cell viability was measured in treated **(A)** nontumorigenic immortal mammary epithelial MCF-10A cells and TNBC **(B)** MDA-MB 231**, (C)** Hs 578t and **(D)** MDA-MB 468 cells. The treatments applied were vehicle addition (paclitaxel, purple), heparan sodium sulfate (50 µM, teal) and OGT 2115 (20 µM, purple-red) or the combination (light blue); heparan sodium sulfate and OGT 2115 were administered for 48 hours, and paclitaxel was added for the final 6 hours to replicate exposure times in patients. The standard deviation was calculated from three independent experiments performed in triplicate. 1-way ANOVA with Tukey’s HSD was applied to ascertain significance. * represents p<0.05 and ** represents p<0.01 when comparing vehicle addition to heparan sodium sulfate, OGT 2115 or the combination.

#### Measurement of heparanase and heparan sulfate expressions in human breast cancer samples by immunohistochemistry

An AMSBIO breast cancer tissue array (120 cores specifically with 82 TNBC cores), two normal breast tissue slides and three ductal carcinoma in situ DCIS tissue slides were stained with heparanase. (Figure 4B) and statistical analysis was performed (Figure 5). The expression of heparanase was compared amongst TNBCs, ER+/PR+, HER2+ breast cancer, normal breast tissue and DCIS. Kruskal-Wallis test showed that there was no significant difference in the percentages of cells that stained at any level, weakly, moderately, strongly positively or negatively for heparanase in tissue sections of normal breast tissue, DCIS and invasive breast cancers. Also, the Kruskal-Wallis test indicated that there was no significant difference in the percentages of cells that stained positively for heparanase in tissue sections of different TNBC breast cancer stages, amongst grades of TNBCs and normal breast tissue and comparing cancers with differing Ki67 expression levels (Supplemental Figures 12-13).

As previously mentioned, heparanase binds to the cell surface and extracellular matrix heparan sulfate chains, and, hence, its expression level may be dependent on heparan sulfate expression levels [33]. However, given that the expression levels are semi-quantitatively assessed by immunohistochemistry, it is difficult to normalize heparanase to heparan sulfate levels. The positive correlation and dependence of heparan expression on heparan sulfate expression levels may account for the paradoxical lack of increased expression of heparanase in tissue sections of invasive cancer compared to those of normal breast tissue and DCIS.

### Effect of heparanase inhibitor on cell viability and eATP

We next analyzed the effects of the combination of heparanase inhibitor (OGT 2115) with chemotherapy (paclitaxel) to assess its impact on the effectiveness of chemotherapy in TNBC MDA-MB 231, Hs 578t and MDA-MB 468 and, in comparison, to non-tumorigenic immortal epithelial mammary MCF-10A cells. For these experiments, all the cell lines were treated with paclitaxel for the final six hours of a 48-hour time course to simulate the duration of systemic exposure to paclitaxel in patients and with OGT 2115 and heparan sodium sulfate for 48 hours (Figure 6). For this reason, we did not see differences in the loss of viability of cells treated with paclitaxel alone (vehicle addition). Yet, in all three TNBC cell lines MDA-MB 231, Hs 578t and MDA-MB 468 when paclitaxel (100 µM) was combined with the heparanase inhibitor OGT 2115 (20 µM), there was a significantly increased percentage loss of cell viability when compared to vehicle addition (paclitaxel alone). Additionally, the percentage loss of cell viability was further significantly enhanced when heparan sodium sulfate was added to the combination of paclitaxel and OGT 2115 when compared to vehicle addition. However, there was no significant change in the percentage loss of viability of MCF-10A cells treated with paclitaxel, OGT 2115 and heparan sodium sulfate.

Under the same conditions, we also assessed the amount of eATP in the supernatants of chemotherapy-treated (paclitaxel-treated) cells (Figure 7). In the presence of OGT 2115 and paclitaxel, we saw significant increases in eATP levels when compared to vehicle addition (paclitaxel alone) in all cell lines immortal MCF-10A cells and TNBC MDA-MB 231, Hs 578t and MDA-MB 468. Moreover, there was an even greater significant increase in eATP with the addition of heparan sodium sulfate to the combination of paclitaxel and OGT 2115 when compared to the vehicle addition. Thus, the heparanase inhibitor OGT 2115 significantly increased eATP release upon chemotherapy treatment and sensitized TNBCs to chemotherapy.

**Figure 7:**
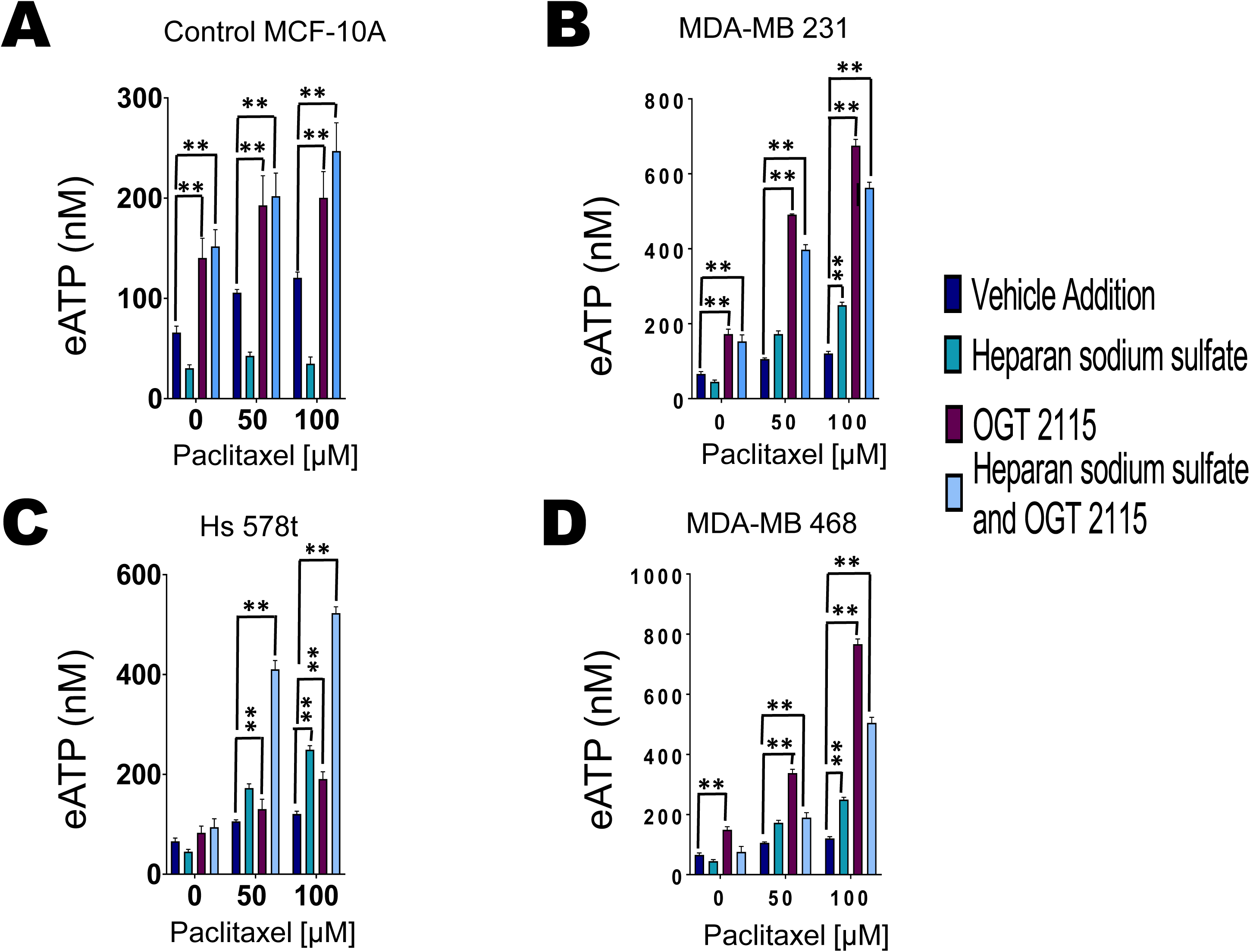
Heparanase inhibitor OGT 2115 and chemotherapeutic agent paclitaxel influence extracellular ATP concentrations. Extracellular ATP concentrations were measured in the supernatants of treated **(A)** nontumorigenic immortal mammary epithelial MCF-10A cells and TNBC **(B)** MDA-MB 231**, (C)** Hs 578t and **(D)** MDA-MB 468 cells. The treatments: vehicle addition (paclitaxel, purple), heparan sodium sulfate (50 µM, teal) and OGT 2115 (20 µM, purple-red), or the combination regimen (light blue); heparan sodium sulfate and OGT 2115 were administered for 48 hours and paclitaxel was added for the final 6 hours to replicate exposure times in patients. The standard deviation was calculated from three independent experiments performed in triplicate. 1-way ANOVA with Tukey’s HSD was applied to ascertain significance. * represents p<0.05 and ** represents p<0.01 when comparing vehicle addition to heparan sodium sulfate, OGT 2115 or the combination regimen.

#### Verification that OGT 2115 inhibits heparanase at the concentration utilized

MDA-MB 468 cell lysates were treated with heparanase, heparanase inhibitor OGT 2115 or their combination; a negative control (no added heparanase or inhibitor) and positive control of heparin (known heparanase inhibitor) were also included. We focused on the syndecan-1 (a protein known to be physiologically post-translationally modified by heparan sulfate) band with a molecular weight at about 50 kDA through immunoblotting where we saw a decrease in expression in the presence of heparanase in contrast to the negative control (Supplemental Figure 14). Enhanced intensity of this band in the presence of OGT 2115 confirms that OGT 2115 is a heparanase inhibitor at the utilized concentration.

### Role of purinergic signaling in the enhancement of chemotherapy-induced TNBC cell death by the application of heparanase inhibitors

We had previously demonstrated that eATP exerts cytotoxic effects on TNBC cells through P2RX4 and P2RX7 receptors [9]. We sought to verify if the exaggerated loss of cell viability in the presence of OGT 2115 is dependent on eATP-induced activation of P2RX4 or P2RX7 (Figure 8). We chose Hs 578t cells for this experiment because we detected the largest increase in eATP and percentage loss of cell viability when this cell line was exposed to the regimen of paclitaxel, OGT 2115 and heparan sodium sulfate. We did observe a significant reversal of the effects of OGT 2115 on cell viability and eATP release upon conducting the experiment in the presence of the P2RX7 inhibitor A438079 (Figures 8A and C) or the P2RX4 inhibitor 5-BDBD (Figures 8B and D). We observed a significant decrease in eATP (p<0.0001) and increased cell viability (p<0.0001) when comparing the combination of paclitaxel with OGT 2115 to that of paclitaxel with OGT 2115 and A43709. We observed a significant decrease in eATP (p<0.0001) and increased cell viability (p<0.0001) when comparing the combination of paclitaxel, heparan sodium sulfate and OGT 2115 to paclitaxel, heparan sodium sulfate, OGT 2115 and A43709. Hence, the P2RX7 blocker A43709 reversed the capacity of OGT 2115, and that of heparan sulfate combined with OGT 2115 to enhance the efficacy and augment eATP release induced by paclitaxel treatment of TNBC cell lines.

**Figure 8:**
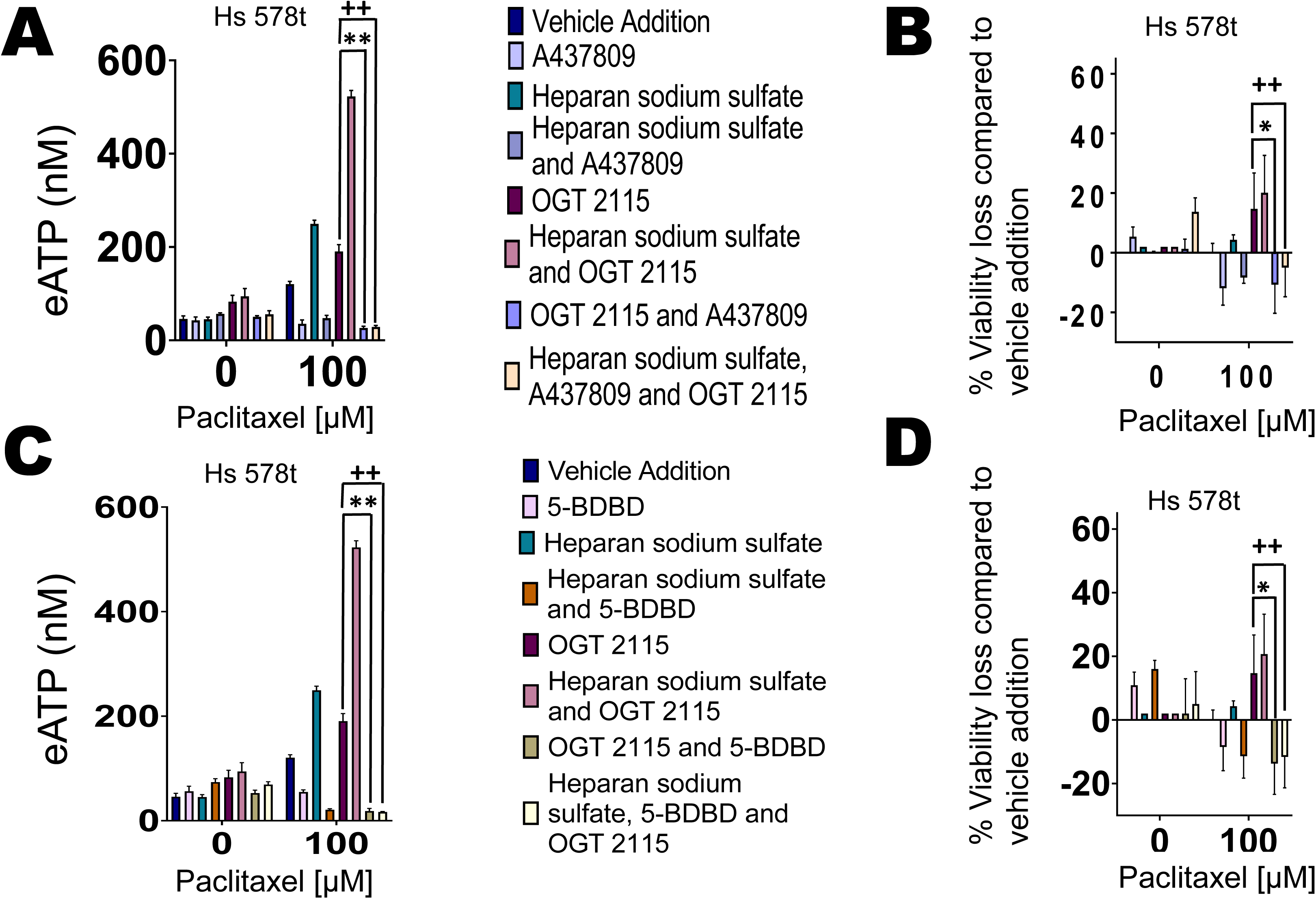
Reversal of heparanase inhibitor’s effects by P2RX4 and P2RX7 inhibitors. For **(A)** and **(B)** Hs 578t cells were treated with OGT 2115 (20 µM, 48 hours), paclitaxel (100 µM, the final 6 hours of the 48-hour time course to replicate exposure times in patients), heparan sodium sulfate (50 µM, 48 hours), A437809 (20 µM, 6 hours), or combination of the different drug agents. The standard deviation was calculated from three independent experiments performed in triplicate. 1-way ANOVA with Tukey’s HSD was applied to ascertain significance. * represents p<0.05 and ** represents p<0.01 when comparing OGT 2115 to the vehicle (paclitaxel), OGT 2115 and A437809 and ++and represents p<0.01 when comparing OGT 2115 to the vehicle (paclitaxel), OGT 2115, A437809 and heparan sodium sulfate. For **(C)** and **(D)** Hs 578t cells were treated with OGT 2115 (20 µM, 48 hours), paclitaxel (100 µM, final 6 hours of the 48-hour time course to replicate exposure times in patients), heparan sodium sulfate (50 µM, 48 hours), 5-BDBD (20 µM, 6 hours), or combinations. The standard deviation was calculated from three independent experiments performed in triplicate. 1-way ANOVA with Tukey’s HSD was applied to ascertain significance. * represents p<0.05 and ** represents p<0.01 when comparing OGT 2115 to a vehicle (paclitaxel), OGT 2115 and 5-BDBD and ++and represents p<0.01 when comparing OGT 2115 to the vehicle (paclitaxel), OGT 2115, 5-BDBD and heparan sodium sulfate.

We observed a significant decrease in eATP (p<0.0001) and increased cell viability (p<0.0001) when comparing the combination of paclitaxel and OGT 2115 to paclitaxel, OGT 2115 and 5-BDBD. We observed a significant decrease in eATP (p<0.0001) and increased cell viability (p<0.0001) when comparing the combination of paclitaxel, heparan sodium sulfate and OGT 2115 to that of paclitaxel, heparan sodium sulfate, OGT 2115 and 5-BDBD. Therefore, 5-BDBD reversed the capacity of OGT 2115 and the combination of heparan sodium sulfate and OGT 2115 to enhance the loss of cell viability induced by paclitaxel by augmenting extracellular ATP release.

These data demonstrate that the exaggerated loss of cell viability observed when OGT 2115 and heparan sulfate are combined with paclitaxel is dependent on the activation of both P2RX4 and P2RX7 by eATP.

### Impact of purinergic signaling on cancer-initiating cells

Additionally, we wanted to evaluate the impact of the heparanase inhibitor OGT 2115 and chemotherapeutic agent paclitaxel on the cancer-initiating cell fraction. Thus, flow cytometry analysis was performed on TNBC cells lines MDA-MB 231, MDA-MB 468 and Hs 578t treated with paclitaxel (100 µM) for the final 6 hours of a 48-hour time course (to stimulate paclitaxel pharmacokinetics when paclitaxel is administered to patients) and OGT 2115 (20 µM) for 48 hours to determine the number of cancer-initiating cells (cells that express high levels of aldehyde dehydrogenase (ALDH) and CD44 but do not express CD24). Paclitaxel enhanced the number of cancer-initiating cells (Figures 9A-C). However, the combination of paclitaxel and OGT 2115 also increased the fraction of ALDH^Hi^CD44+CD24 (cancer-initiating cells) in this assay.

**Figure 9:**
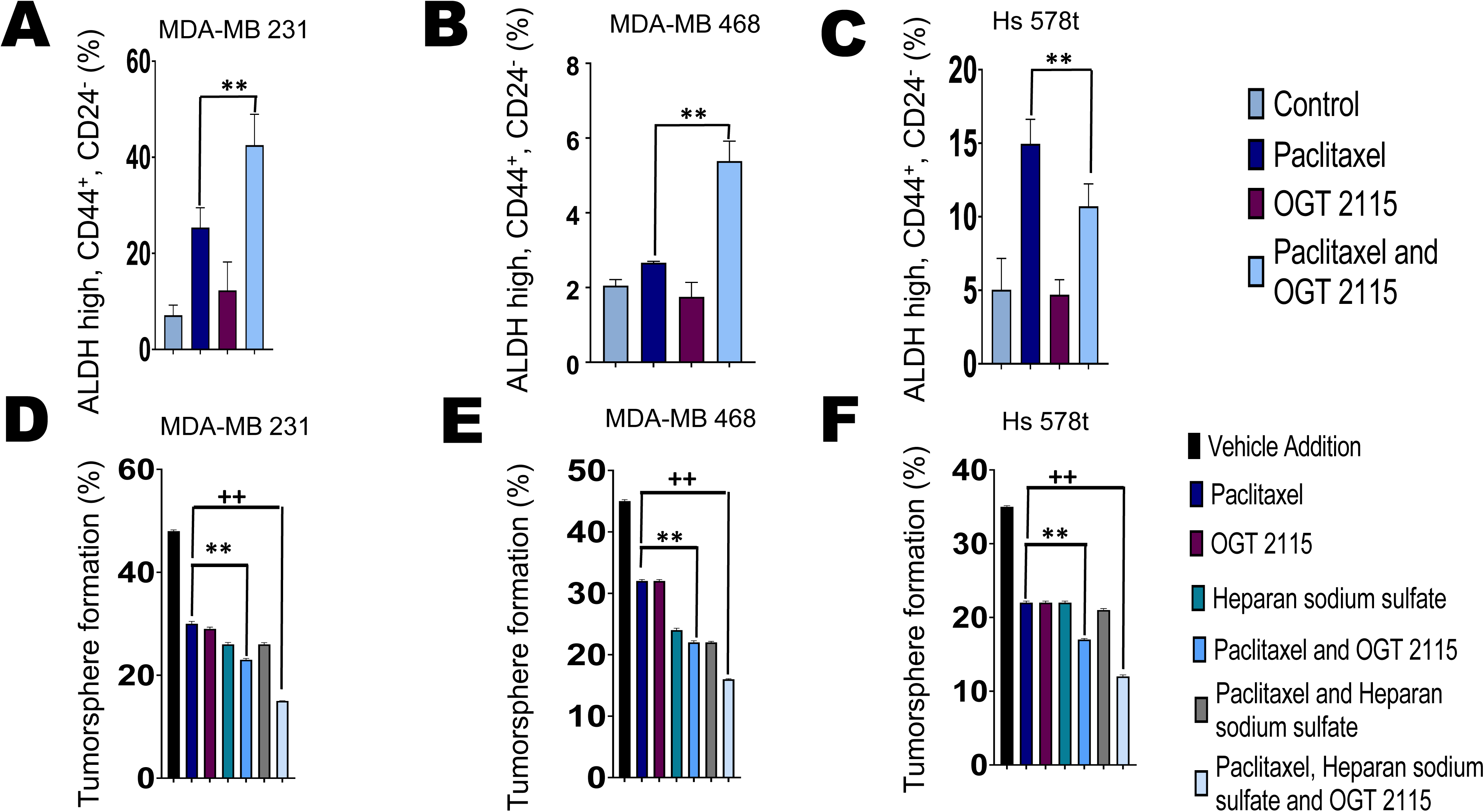
Effect of heparanase inhibitor OGT 2115 combined with chemotherapy on cancer-initiating cell fraction of treated triple-negative breast cancer cells. The three TNBC cell lines **(A)** MDA-MB 231, **(B)** MDA-MB 468 and **(C)** Hs 578t were treated with OGT 2115 (20 µM, 48 hours) and/or paclitaxel (100 µM, final 6 hours of the 48-hour time course to replicate exposure times in patients). Paclitaxel increased the number of cancer-initiating cells (cells that express high levels of ALDH and CD44 but do not express CD24). However, the combination of paclitaxel and OGT 2115 also increased the number of cancer-initiating cells. 1-way ANOVA with Tukey’s HSD was applied to ascertain significance. * represents p<0.05 and ** represents p<0.01 when comparing paclitaxel to paclitaxel and OGT 2115. Effects on cancer-initiating cells were also determined through the tumorsphere formation efficiency assay in which triple-negative breast cancer cell lines **(D)** MDA-MB 231 **(E)** MDA-MB 468 and **(F)** Hs 578t cells were treated with vehicle (DMSO), paclitaxel (100 µM, final 6 hours of the 48-hour time course to replicate exposure times in patients), heparan sodium sulfate (50 µM, 48 hours), OGT 2115 (20 µM, 48 hours) or the different combinations listed. Treated triple-negative breast cancer cells were washed, passed through cell strainers, collected, and plated at approximately one cell per well on round-bottom low-attachment 96-well plates and tumorspheres were allowed to form for 7 days. The fraction of wells plated with at least one live cell that was positive for tumorspheres after seven days were counted using Etaluma™ Lumascope 620. The combination regimens showed a significant decrease in tumorsphere formation when compared to the single agent treatments of vehicle, paclitaxel, heparan sodium sulfate or OGT 2115 treated cells. 1-way ANOVA with Tukey’s HSD was applied to ascertain significance. ** represents p<0.01 when comparing paclitaxel to paclitaxel and OGT 2115.++ ** represents p<0.01 when comparing paclitaxel to paclitaxel, heparan sodium sulfate and OGT 2115.

We then applied an orthogonal assay to analyze the effects of purinergic signaling on cancer-initiating cells by carrying out the tumorsphere formation efficiency assay. TNBC cell lines were treated with paclitaxel, OGT 2115 and/or heparan sodium sulfate and processed and maintained as spheroids as described in the methods. After seven days, we observed that the fraction of wells that had been plated with at least one viable cell that showed tumorsphere formation was decreased compared to vehicle treatment for all TNBC cells MDA-MB 231, MDA-MB 468 and Hs 578t in the presence of paclitaxel, heparan sodium sulfate and OGT 2115 (Figure 9D-F). Images were also recorded and provided in the supplemental section (Supplemental Figures 22-24). Hence, this data suggests that the heparanase inhibitor OGT 2115 suppresses the cancer-initiating cell fraction.

As the tumorsphere formation efficiency assay is a functional assay of cancer-initiating cell properties, it may provide a more accurate results than flow cytometry, which relies on stem cell markers alone.

## Discussion

Chemotherapy is still the standard treatment for triple-negative breast cancer. A major drawback of chemotherapy is its inability to eliminate metastatic disease, despite transient responses. Therapeutic approaches that expand and increase responses are urgently needed. eATP, in the high micromolar to millimolar range, has been demonstrated to be cytotoxic to cancer cell lines. We have demonstrated that chemotherapy treatment augments eATP release from TNBC cells. We also showed that ecto-ATPase inhibitors enhance chemotherapy-induced eATP release from TNBC cells and augment chemotherapy-induced cell death. However, this strategy is limited by the presence of multiple families of ecto-ATPases in humans, each with multiple members, thus, complicating the design of synthetic inhibitors.

Polysulfated polysaccharides have been revealed to inhibit multiple classes of ecto-ATPases. The endogenous polysulfated polysaccharide heparan sulfate inhibits multiple families of ectoATPases, thus attenuating the degradation of ATP [11, 12].. Therefore, we hypothesized that increasing heparan sulfate in the microenvironment of triple-negative cancer cells using heparanase inhibitors would enhance eATP concentrations in the pericellular environment of chemotherapy-treated TNBC cells and hence, augment chemotherapy-induced cell death.

Consistent with this hypothesis, our heparan sulfate expression results showed that TNBCs expressed significantly less heparan sulfate on the cell surface. Paradoxically, our immunoblot results revealed that heparanase is highly expressed intracellularly and extracellularly in immortal mammary epithelial cells when compared to TNBC cells. Also paradoxically, the ELISA and immunohistochemistry results did not show increased heparanase expression levels in TNBC cell lines or invasive breast cancers as compared to immortal mammary epithelial cells or normal breast tissue and DCIS respectively. As noted in the results, this could be due to the dependence of heparanase on binding to heparan sulfate for its expression and its transcriptional induction by heparan sulfate [33, 34]. Hence, the ratio of heparanase to heparan sulfate expression may be a better measure of heparanase activity. However, this would need further validation with EXT1 and EXT2 (heparan sulfate synthetic enzymes) overexpressing and/or knock out cell lines to confirm that heparanase expression varies positively and in a linear fashion with heparan sulfate expression.

Our lab has already demonstrated that extracellular ATP in the high micromolar to millimolar range of concentrations is toxic to TNBC cells and that inhibitors of each of the major classes of ecto-ATPases augment chemotherapy-induced increases in eATP. One deficiency of this approach as a promising therapeutic strategy is the need to apply various inhibitors for each of the classes of ecto-ATPases to maximally enhance eATP levels. As heparan sulfate inhibits multiple classes of ectoATPases, we sought to determine if heparanase inhibitors, which inhibit heparan sulfate degradation, can augment chemotherapy-induced cytotoxicity and exacerbate chemotherapy-induced eATP release [11, 12].We revealed that the combination of the heparanase inhibitor OGT 2115 and chemotherapy (paclitaxel) increased extracellular eATP concentrations and enhanced the chemotherapeutic response in TNBCs leading to a larger loss of viability. Additionally, the effects of the heparanase inhibitor on eATP levels and TNBC cell death were changed to basal levels by specific inhibitors of P2RX4 and P2RX7 eATP receptors, affirming that these effects are dependent on these purinergic receptors; we have demonstrated before that these receptors are necessary for chemotherapy-induced eATP release from TNBC cells and its cytotoxic effects [9].

Additionally, we evaluated the effects of combinations of heparanase inhibitor and chemotherapy on cancer-initiating cells, as failure to eradicate these cells results in the failure of cytotoxic chemotherapy to eliminate metastatic TNBC [29, 31, 32, 35]. We revealed that in the presence of chemotherapy and the heparanase inhibitor, there were fewer cancer-initiating cells, as assessed by flow cytometry, across all TNBC cell lines.

As eATP is a known immune danger signal and its metabolite adenosine is considered a potent immunosuppressant; hence additional research is needed to assess the immune effects of ecto-ATPase inhibition by heparan sulfate through the use of immunocompetent *in vivo* models of TNBC. Additionally, more work is needed to assess if the cytotoxic effects on TNBC cells occur through non-specific permeabilization of P2RX7 ion-coupled channels or downstream activation of pyroptosis. We will focus our future research on these aspects of heparan sulfate in the tumor microenvironment.

## Conclusion

Heparanase inhibition possesses the ability to sensitize triple-negative breast cancer cell lines to chemotherapy by increasing extracellular ATP concentrations in the microenvironment of chemotherapy-treated cells. Therefore, heparanase inhibitors may have the possibility to generate deeper and more durable responses in combination with chemotherapy. A major focus of our future goals would be to confirm these hypotheses *in vivo*. As extracellular ATP is a noted immune danger signal, it will be critical to establish the immune effects of this therapeutic strategy.

## Declarations

### Ethics approval and consent to participate

Not applicable.

### Consent for publication

Not applicable.

### Availability of data and materials

The datasets used and/or analyzed during the current study are available from the corresponding author upon reasonable request.

### Authors’ contributions

All authors contributed to the review and analysis. JM performed a majority of the assays with LM carrying out the immunohistochemistry analysis. JM and MC conceived of and designed the experiments, reviewed the data, and authored and edited the manuscript.

## Acknowledgments

Not applicable.

