## Supplemental Figures for "The role of heparan sulfate in enhancing the chemotherapeutic response in triple-negative breast cancer"

### Slide 1
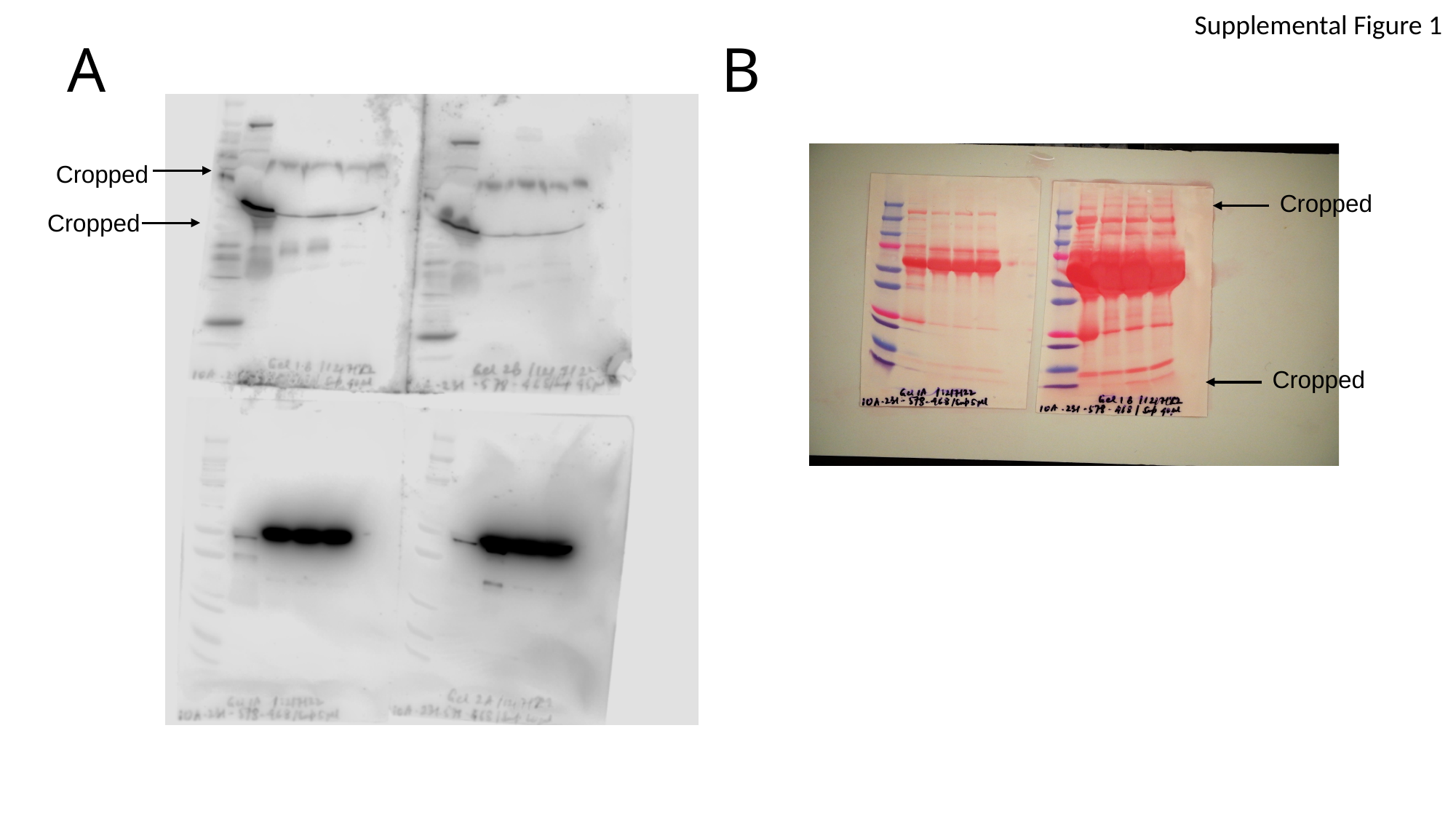

Supplemental Figure 1
A
B
 Cropped
 Cropped
 Cropped
 Cropped

### Slide 2
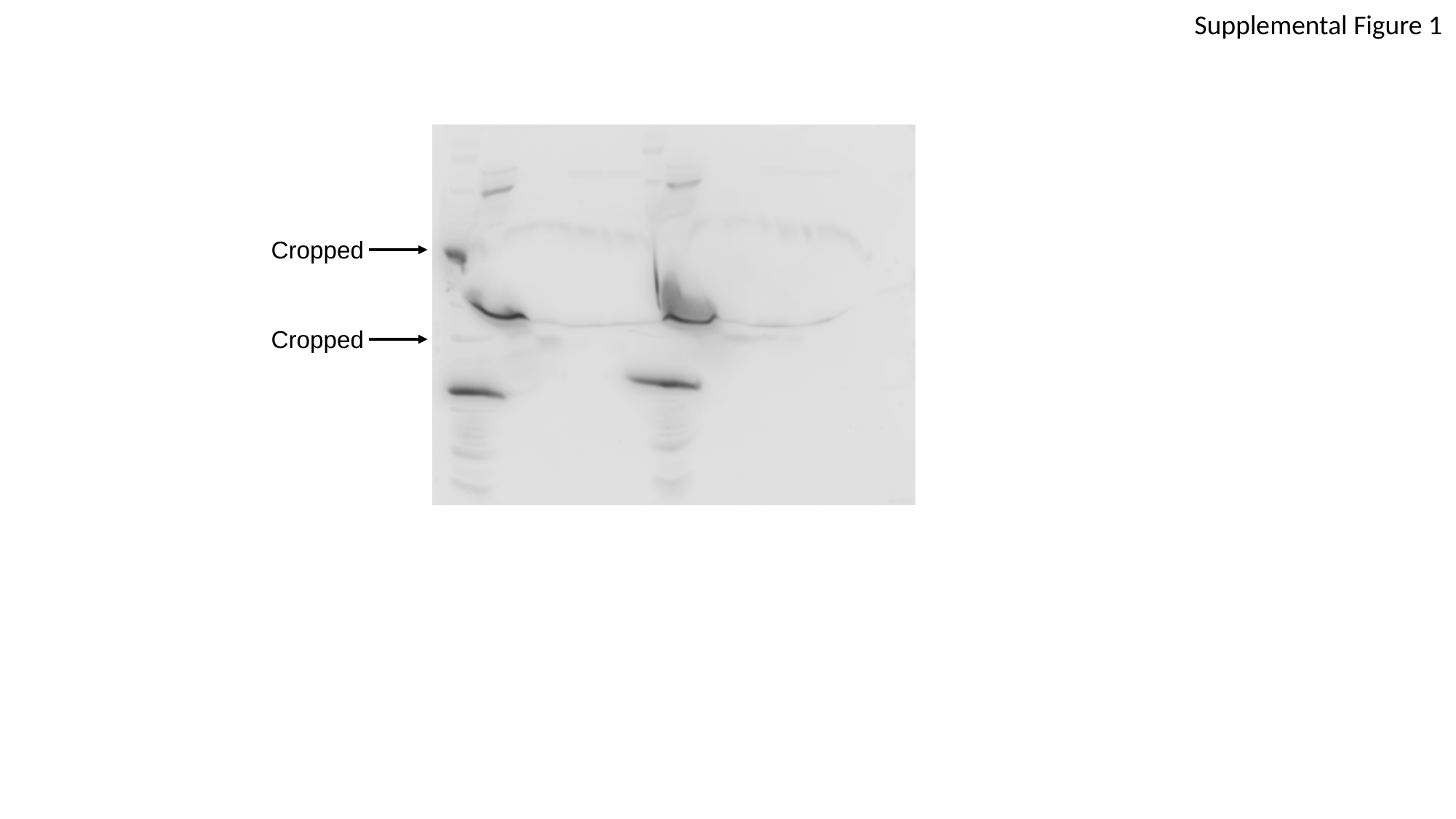

Supplemental Figure 1
 Cropped
 Cropped

### Slide 3
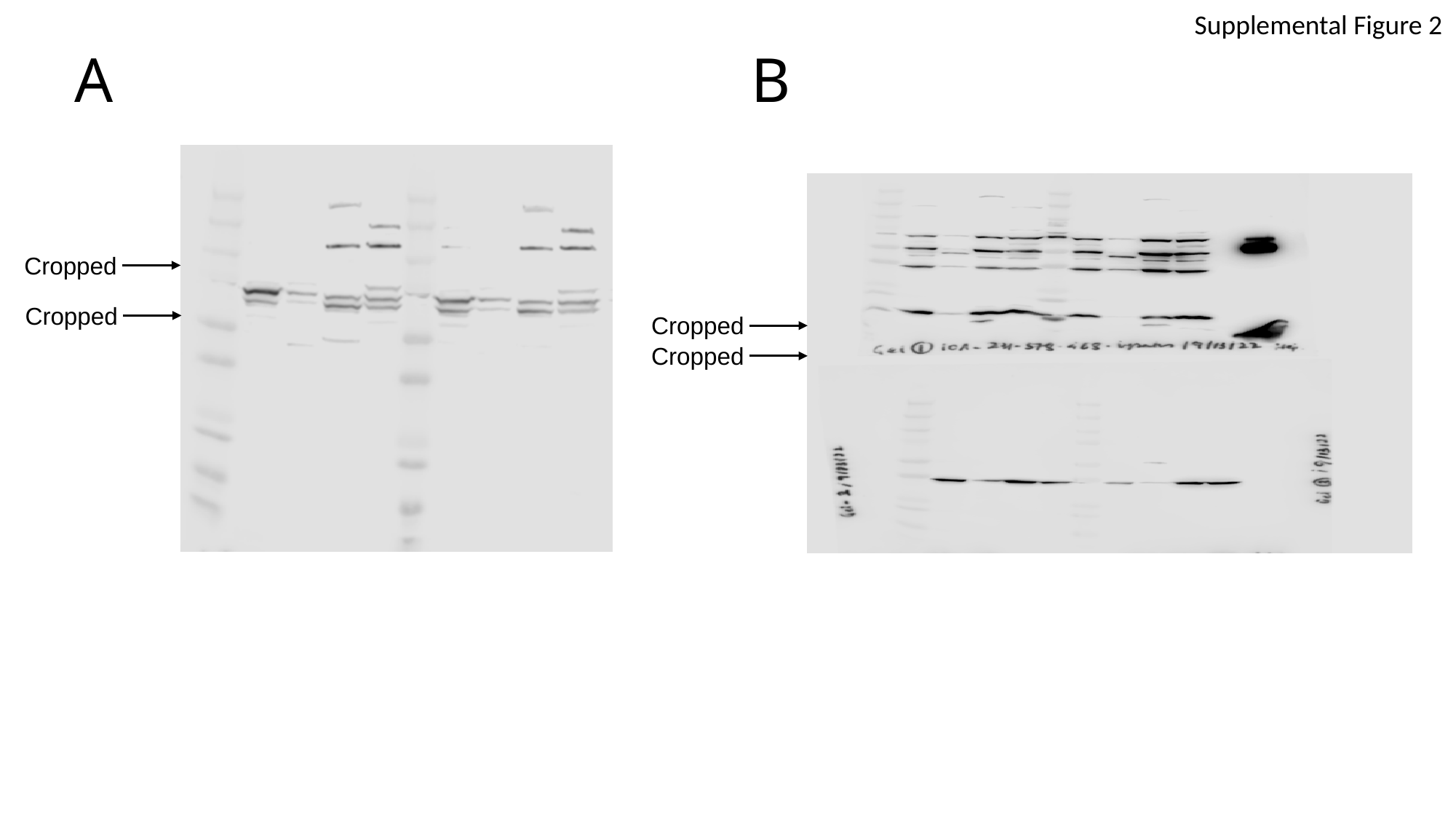

Supplemental Figure 2
A
B
 Cropped
 Cropped
 Cropped
 Cropped

### Slide 4
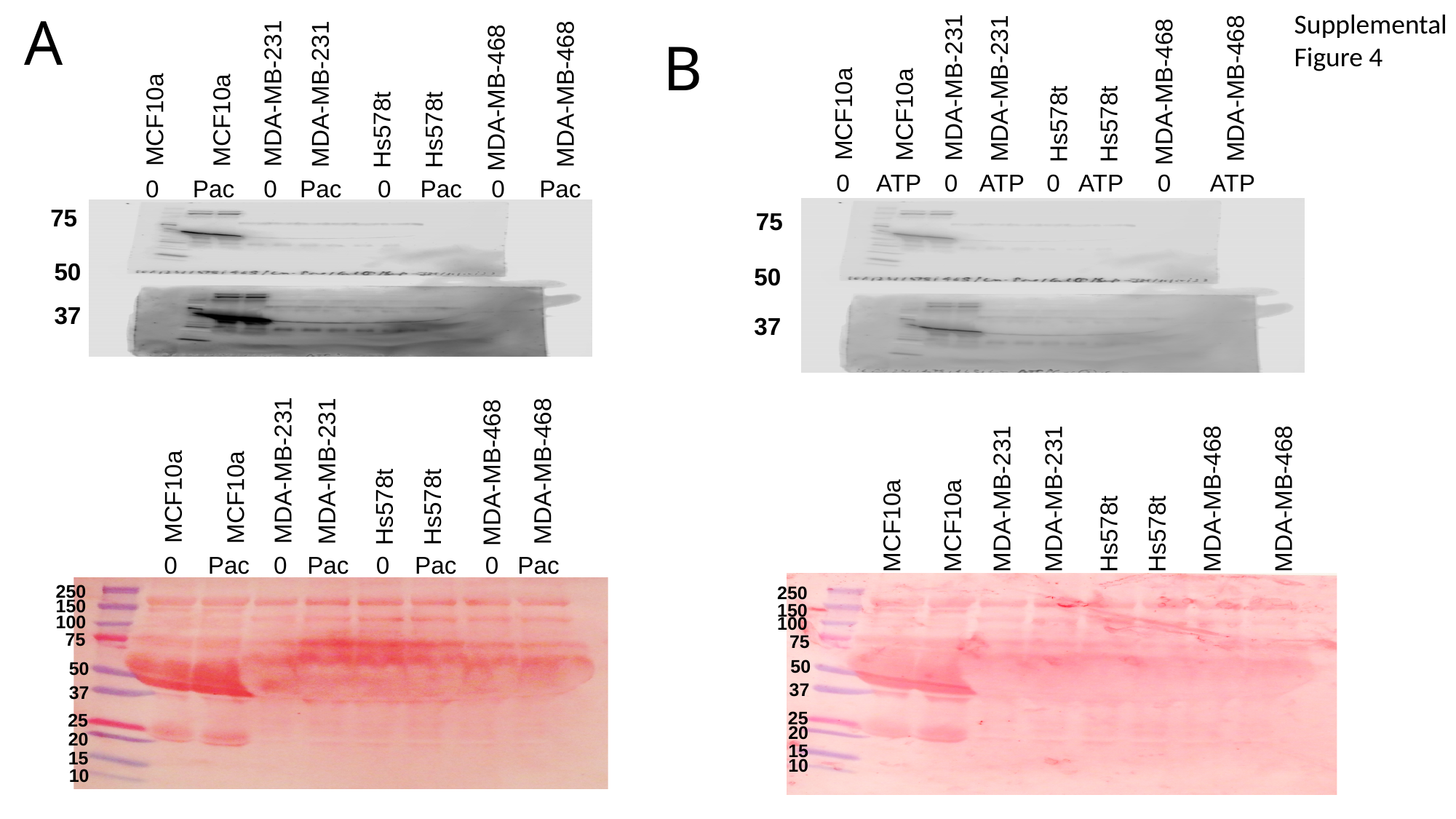

MDA-MB-231
MDA-MB-231
MDA-MB-468
Hs578t
Hs578t
MDA-MB-468
MCF10a
MCF10a
0
ATP
0
ATP
0
ATP
0
ATP
75
50
37
A
Supplemental Figure 4
MDA-MB-231
MDA-MB-231
MDA-MB-468
Hs578t
Hs578t
MDA-MB-468
MCF10a
MCF10a
0
Pac
0
Pac
0
Pac
0
Pac
B
75
50
37
MDA-MB-231
MDA-MB-231
MDA-MB-468
Hs578t
Hs578t
MDA-MB-468
MCF10a
MCF10a
0
Pac
0
Pac
0
Pac
0
Pac
250
150
100
75
50
37
25
20
15
10
MDA-MB-231
MDA-MB-231
Hs578t
Hs578t
MDA-MB-468
MDA-MB-468
MCF10a
MCF10a
0
ATP
0
ATP
0
ATP
0
ATP
250
150
100
75
50
37
25
20
15
10

### Slide 5
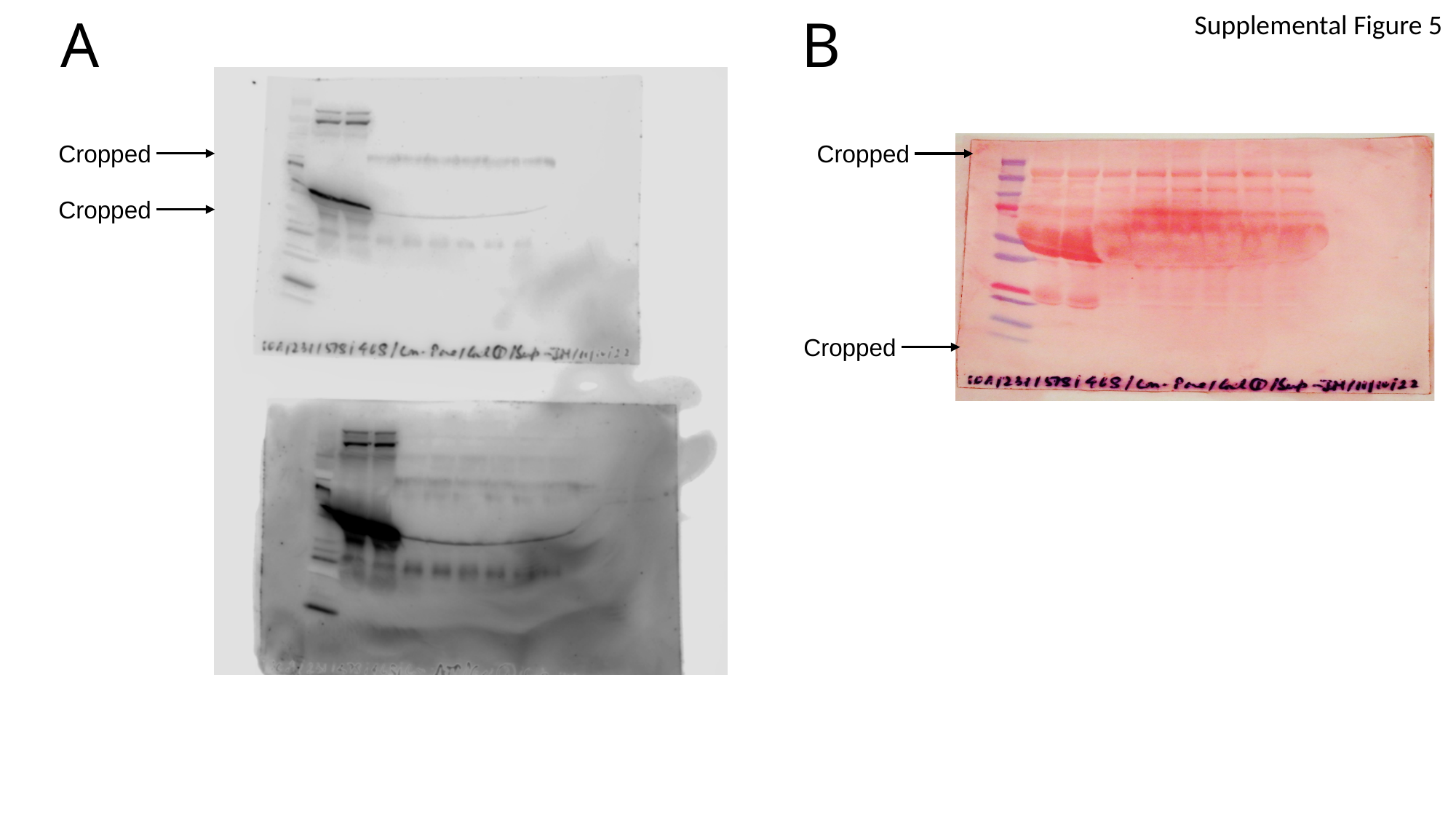

B
A
Supplemental Figure 5
 Cropped
 Cropped
 Cropped
 Cropped

### Slide 6
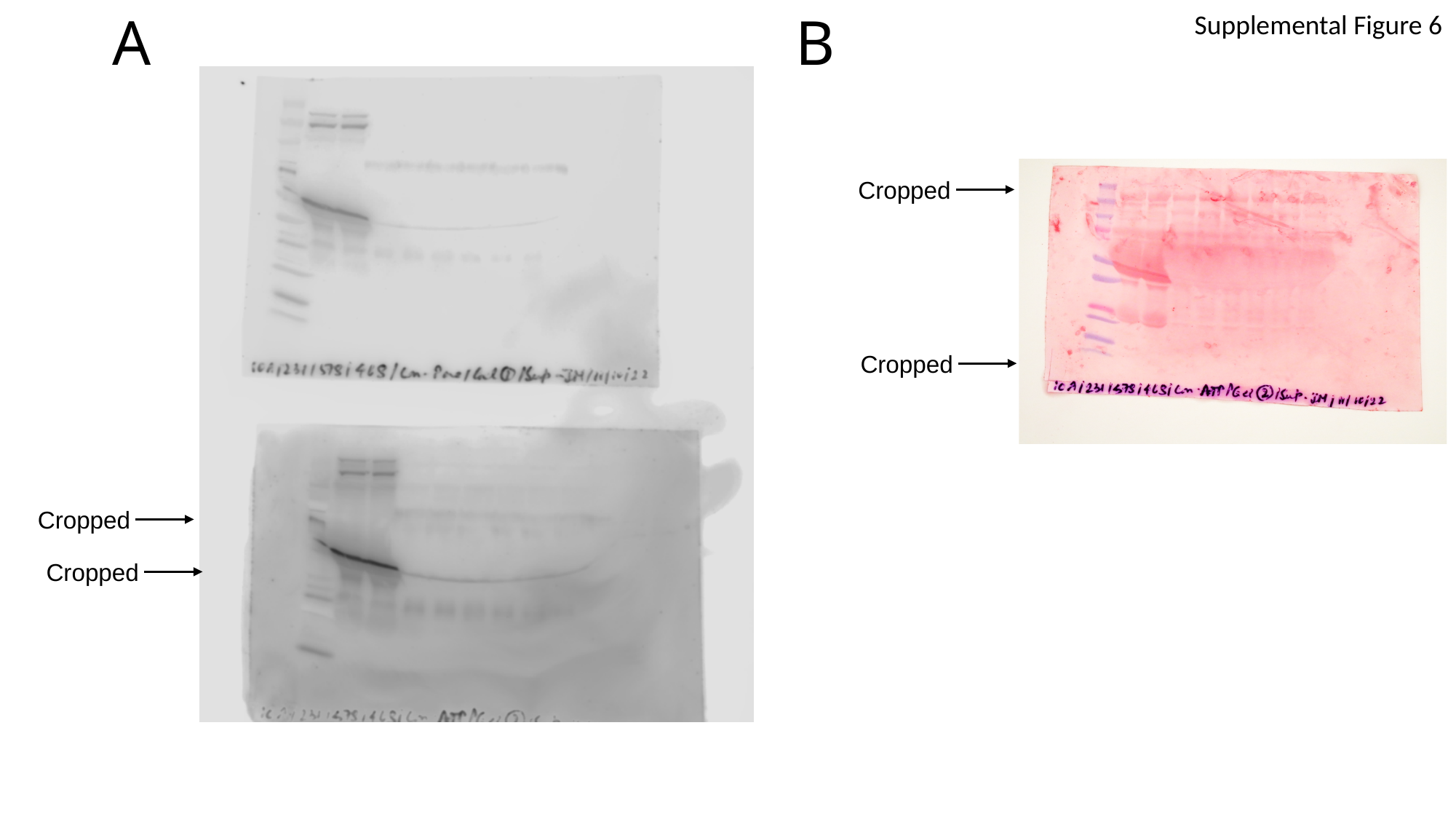

B
A
Supplemental Figure 6
 Cropped
 Cropped
 Cropped
 Cropped

### Slide 7
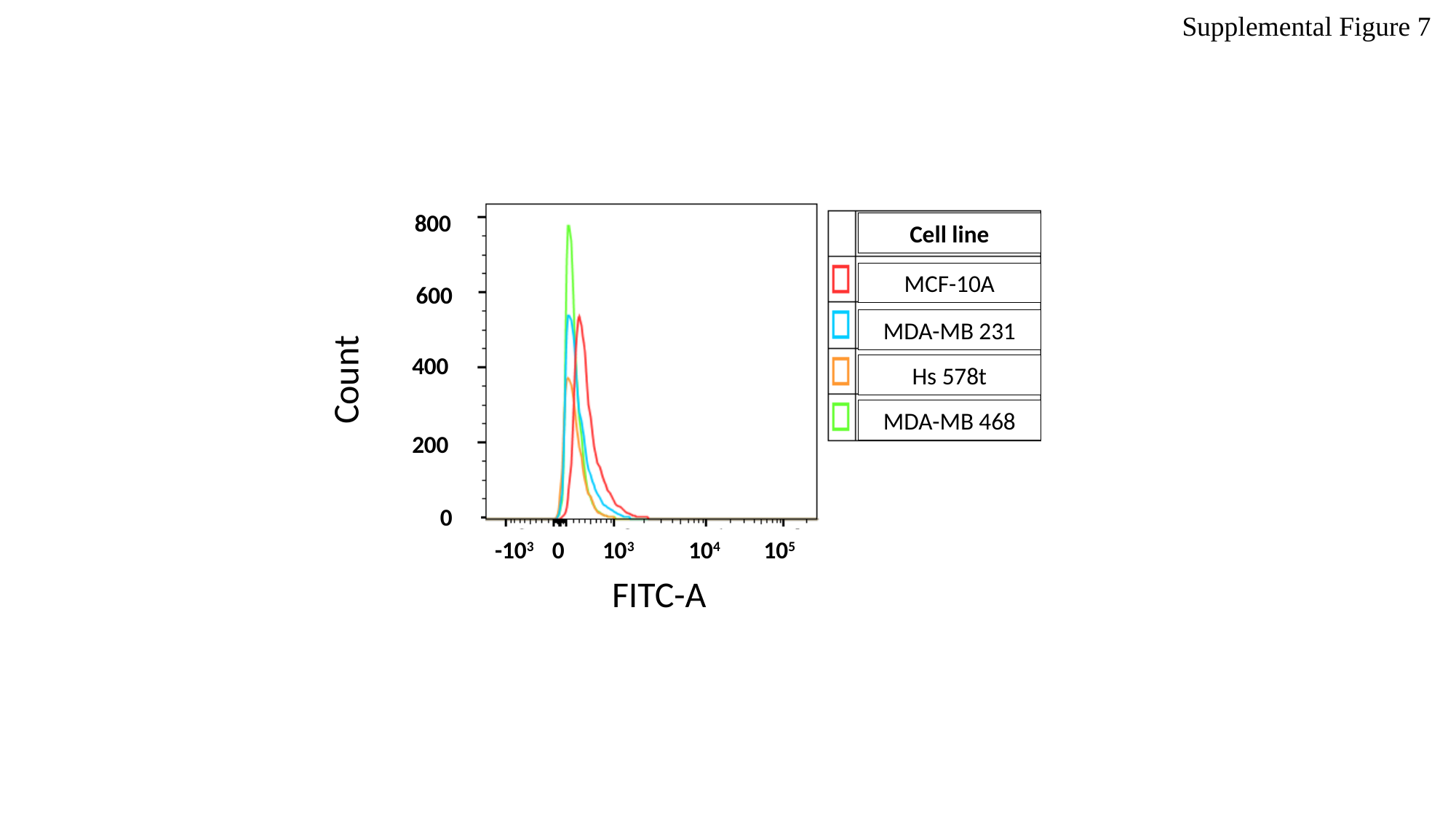

Supplemental Figure 7
800
Cell line
MCF-10A
Count
600
MDA-MB 231
400
Hs 578t
MDA-MB 468
200
0
-10^3^ 0 10^3^ 10^4^ 10^5^
FITC-A

### Slide 8
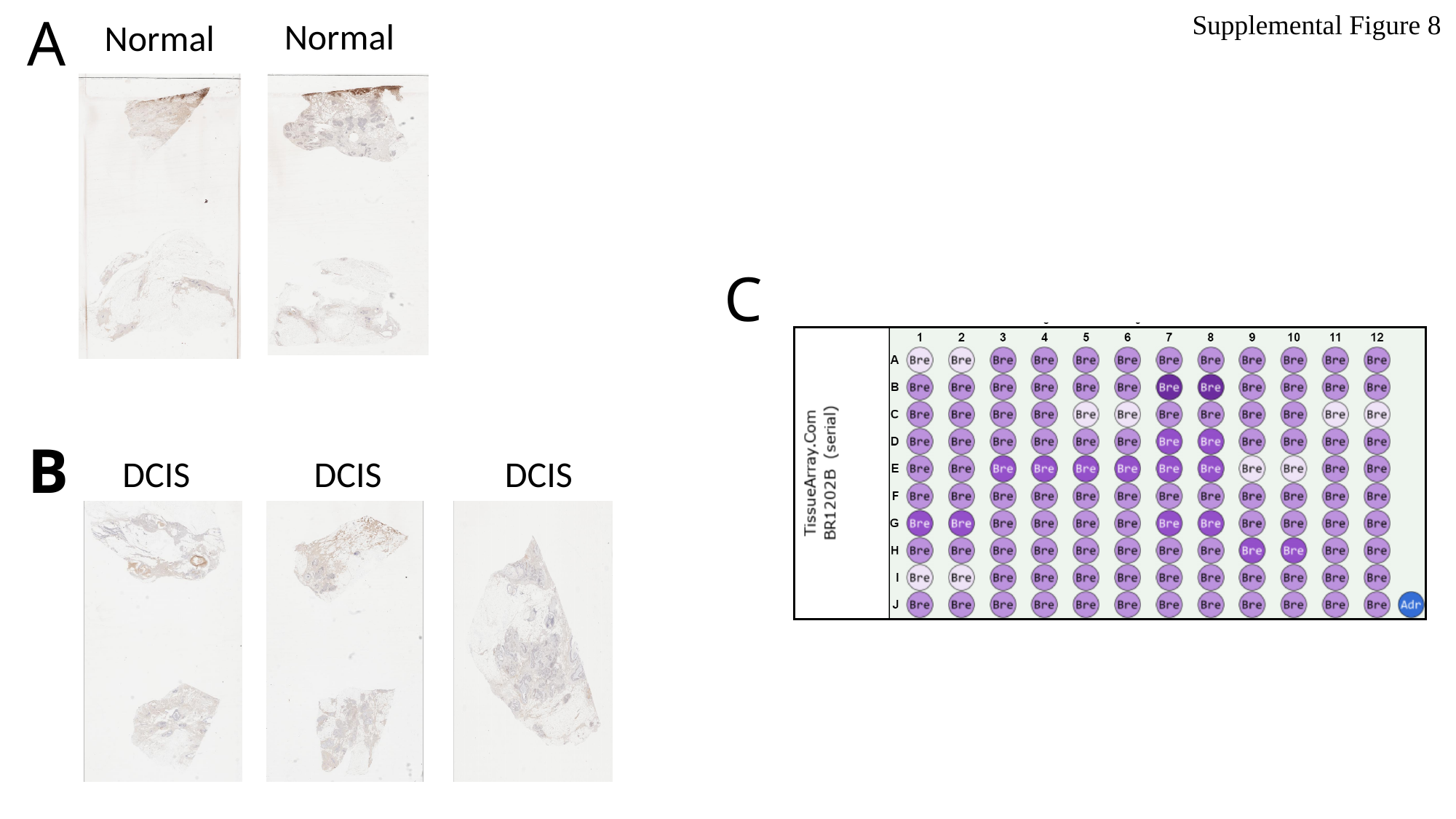

A
Supplemental Figure 8
Normal
Normal
C
B
DCIS
DCIS
DCIS

### Slide 9
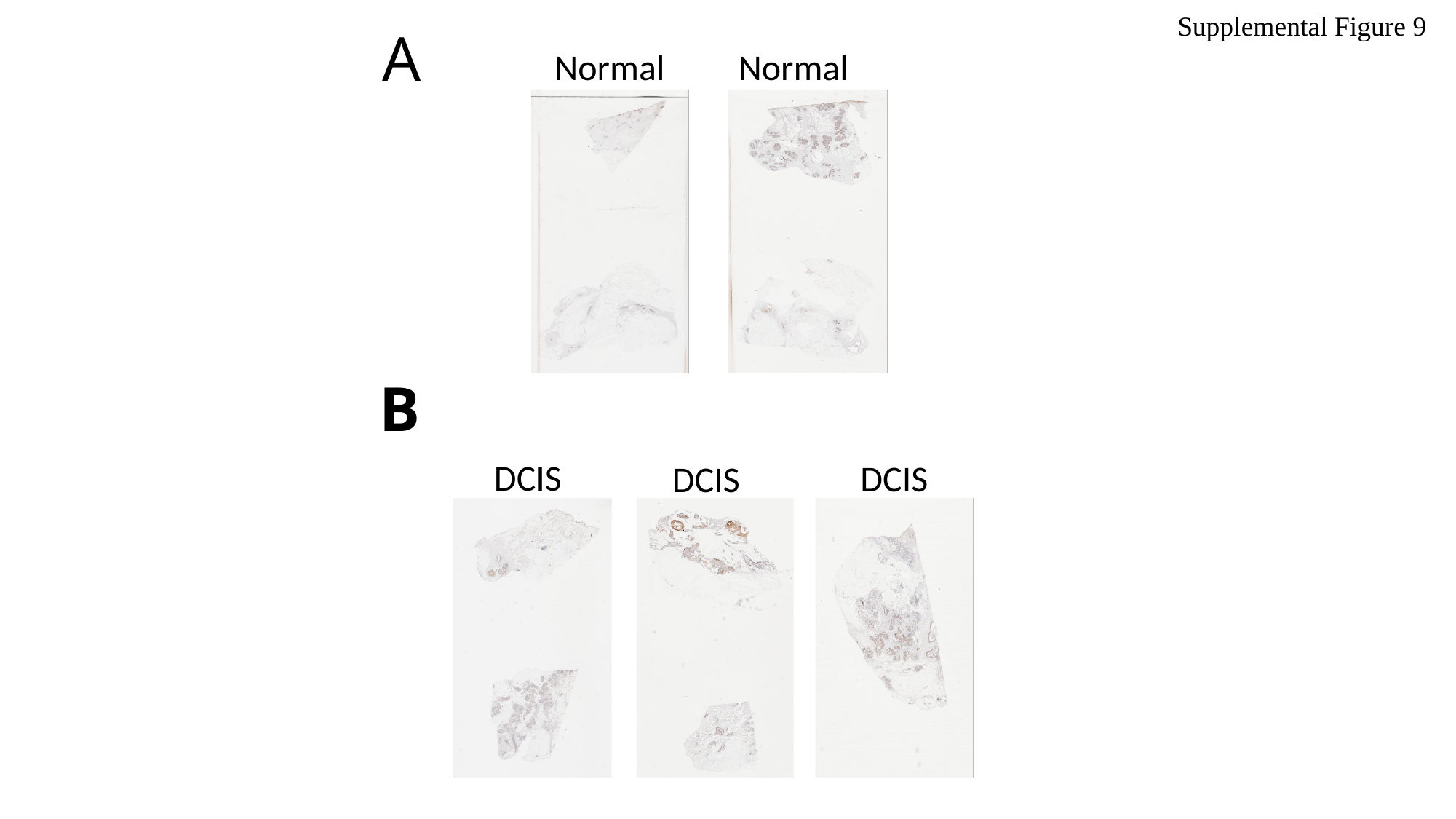

Supplemental Figure 9
A
Normal
Normal
B
DCIS
DCIS
DCIS

### Slide 10
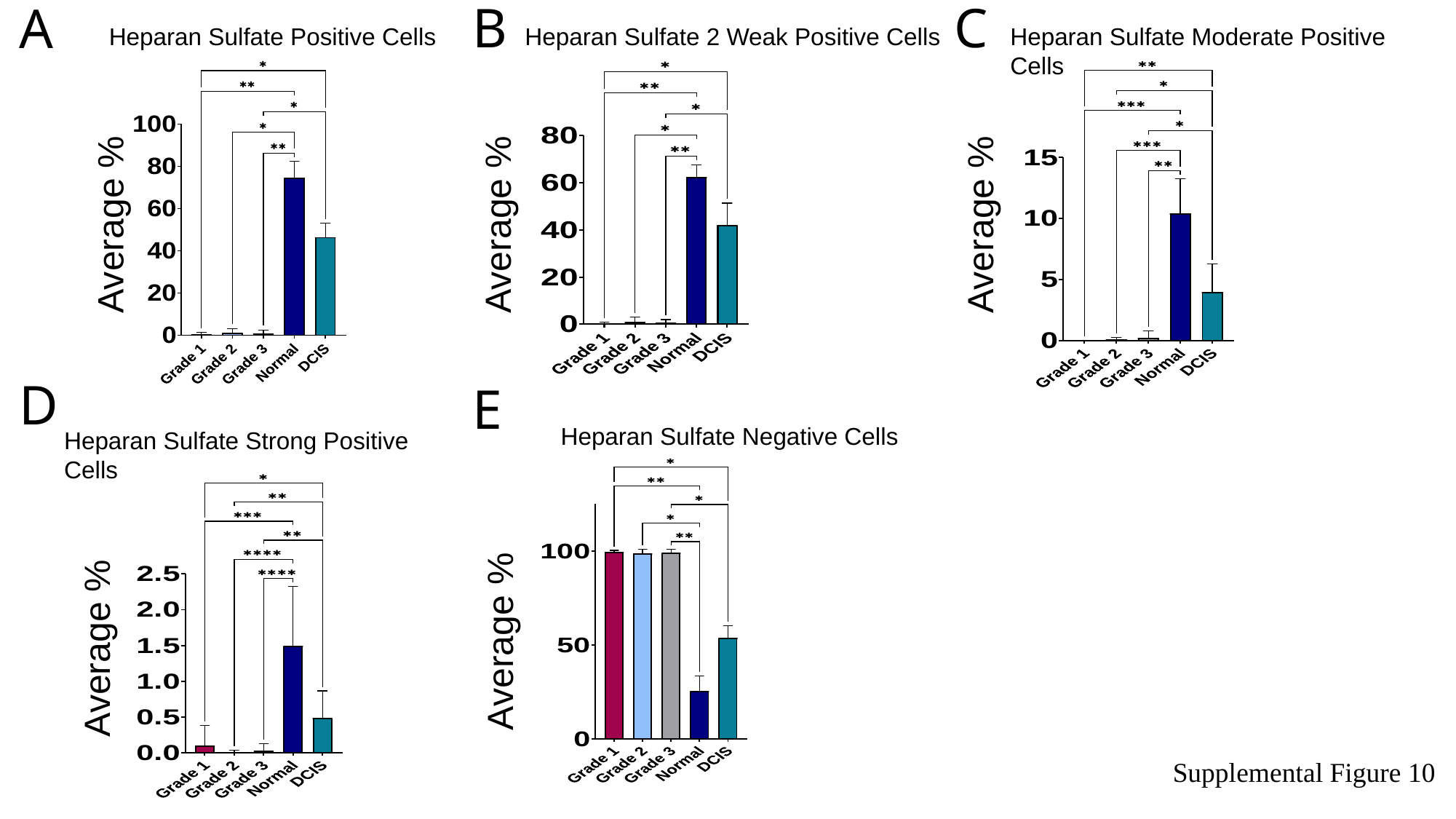

Heparan Sulfate 2 Weak Positive Cells
Heparan Sulfate Moderate Positive Cells
Heparan Sulfate Positive Cells
B
C
A
Average %
Average %
Average %
Heparan Sulfate Strong Positive Cells
Heparan Sulfate Negative Cells
D
E
Average %
Average %
Supplemental Figure 10

### Slide 11
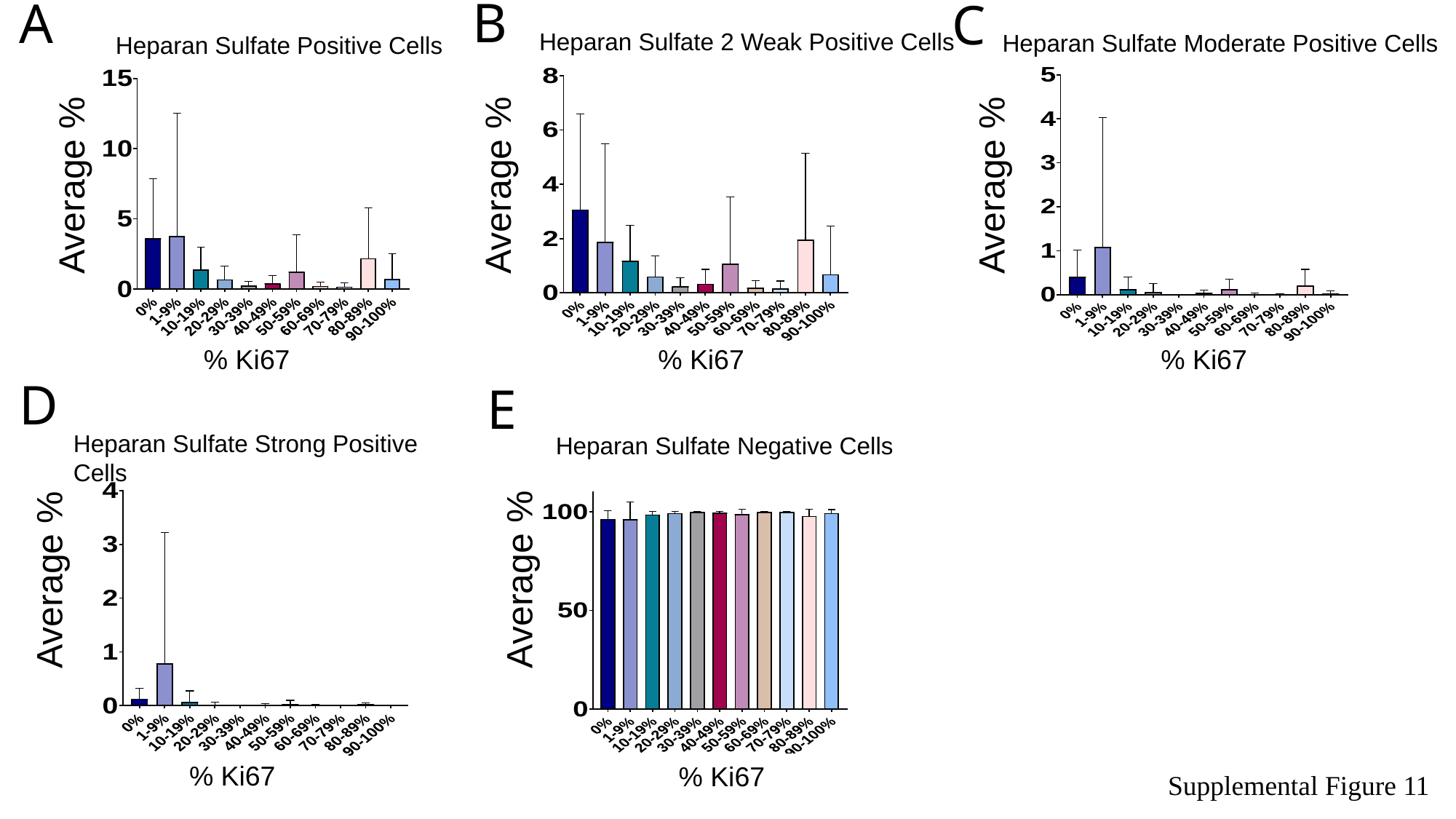

Heparan Sulfate Moderate Positive Cells
Heparan Sulfate 2 Weak Positive Cells
Heparan Sulfate Positive Cells
B
A
C
Average %
Average %
Average %
Heparan Sulfate Strong Positive Cells
Heparan Sulfate Negative Cells
% Ki67
% Ki67
% Ki67
D
E
Average %
Average %
% Ki67
% Ki67
Supplemental Figure 11

### Slide 12
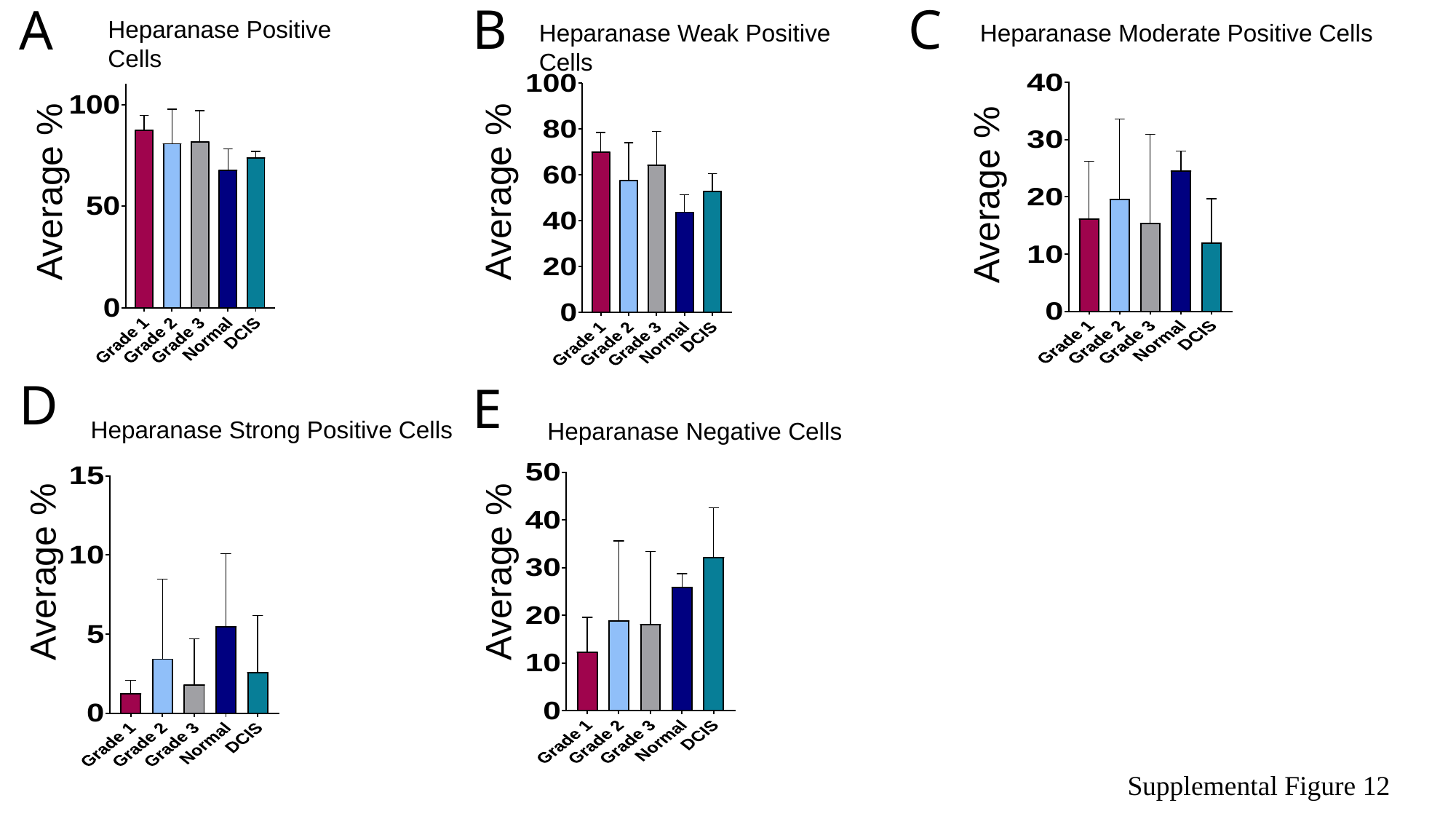

Heparanase Moderate Positive Cells
Heparanase Weak Positive Cells
Heparanase Positive Cells
B
C
A
Average %
Average %
Average %
Heparanase Strong Positive Cells
Heparanase Negative Cells
D
E
Average %
Average %
Supplemental Figure 12

### Slide 13
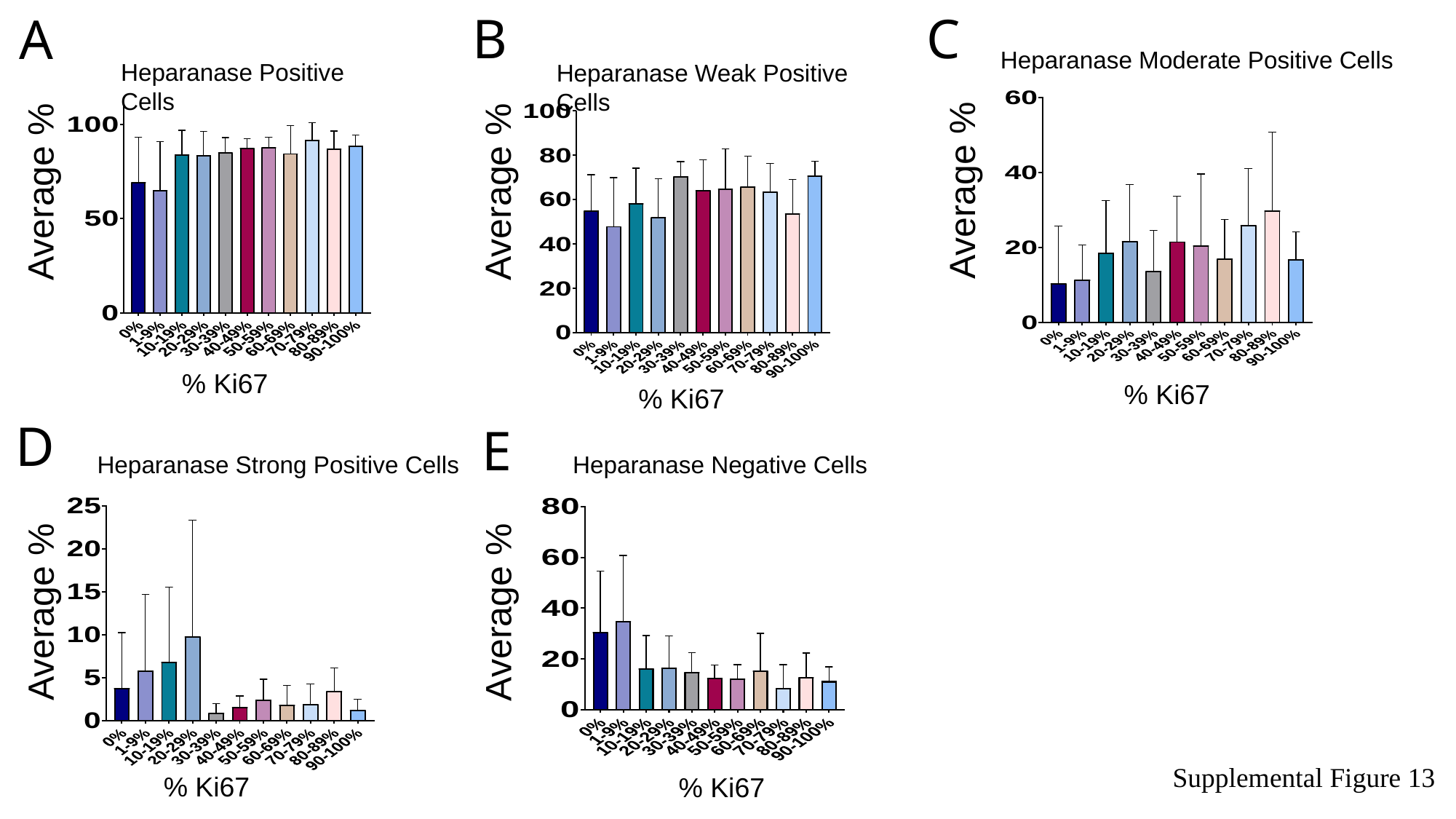

Heparanase Moderate Positive Cells
Heparanase Weak Positive Cells
Heparanase Positive Cells
B
C
A
Average %
Average %
Average %
Heparanase Strong Positive Cells
Heparanase Negative Cells
% Ki67
% Ki67
% Ki67
D
E
Average %
Average %
% Ki67
% Ki67
Supplemental Figure 13

### Slide 14
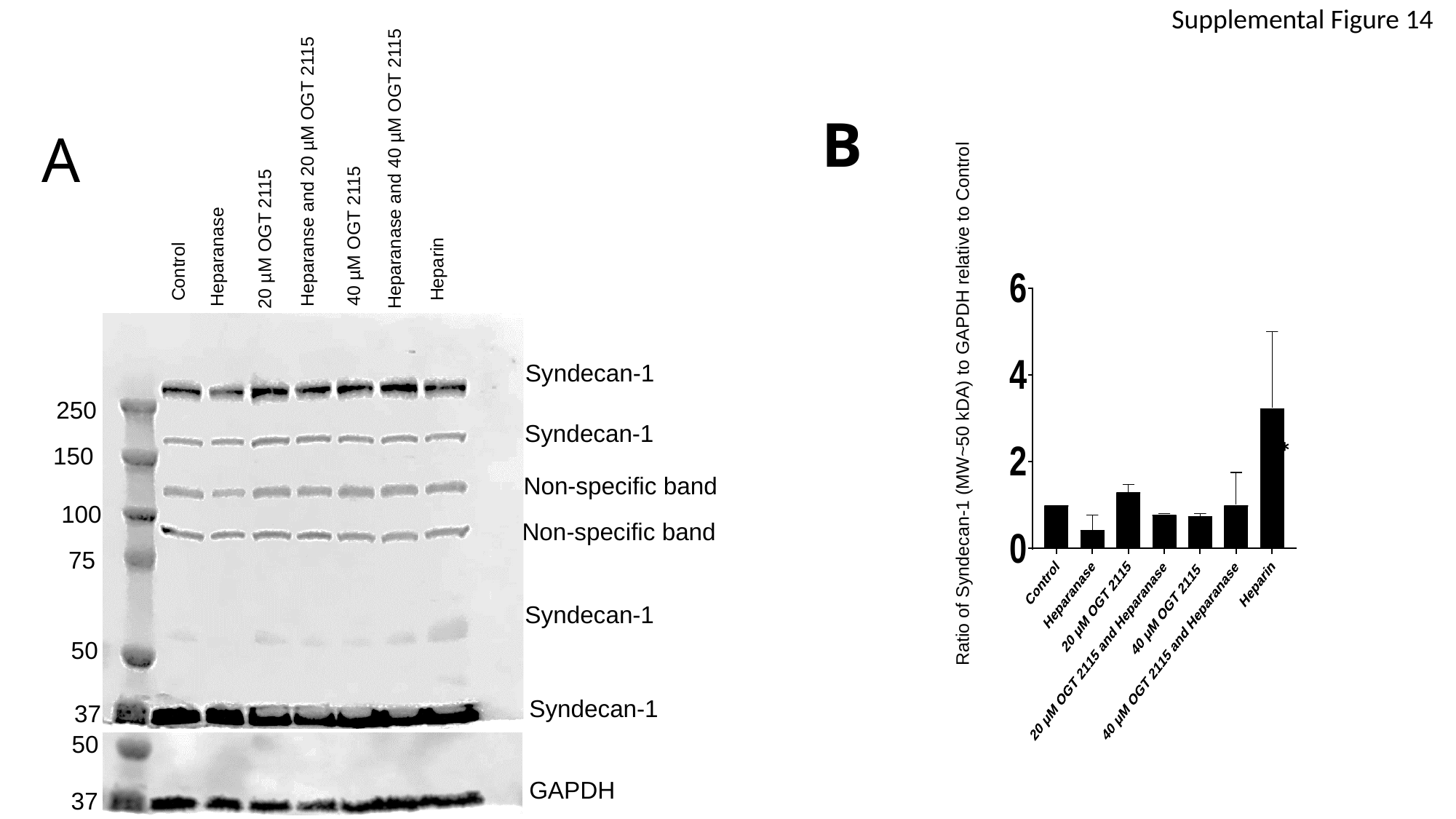

Heparanase and 40 µM OGT 2115
Supplemental Figure 14
Heparanse and 20 µM OGT 2115
Ratio of Syndecan-1 (MW~50 kDA) to GAPDH relative to Control
B
A
20 µM OGT 2115
40 µM OGT 2115
Heparanase
Control
Heparin
Syndecan-1
250
Syndecan-1
*
150
Non-specific band
100
Non-specific band
75
Syndecan-1
50
Syndecan-1
37
50
GAPDH
37

### Slide 15
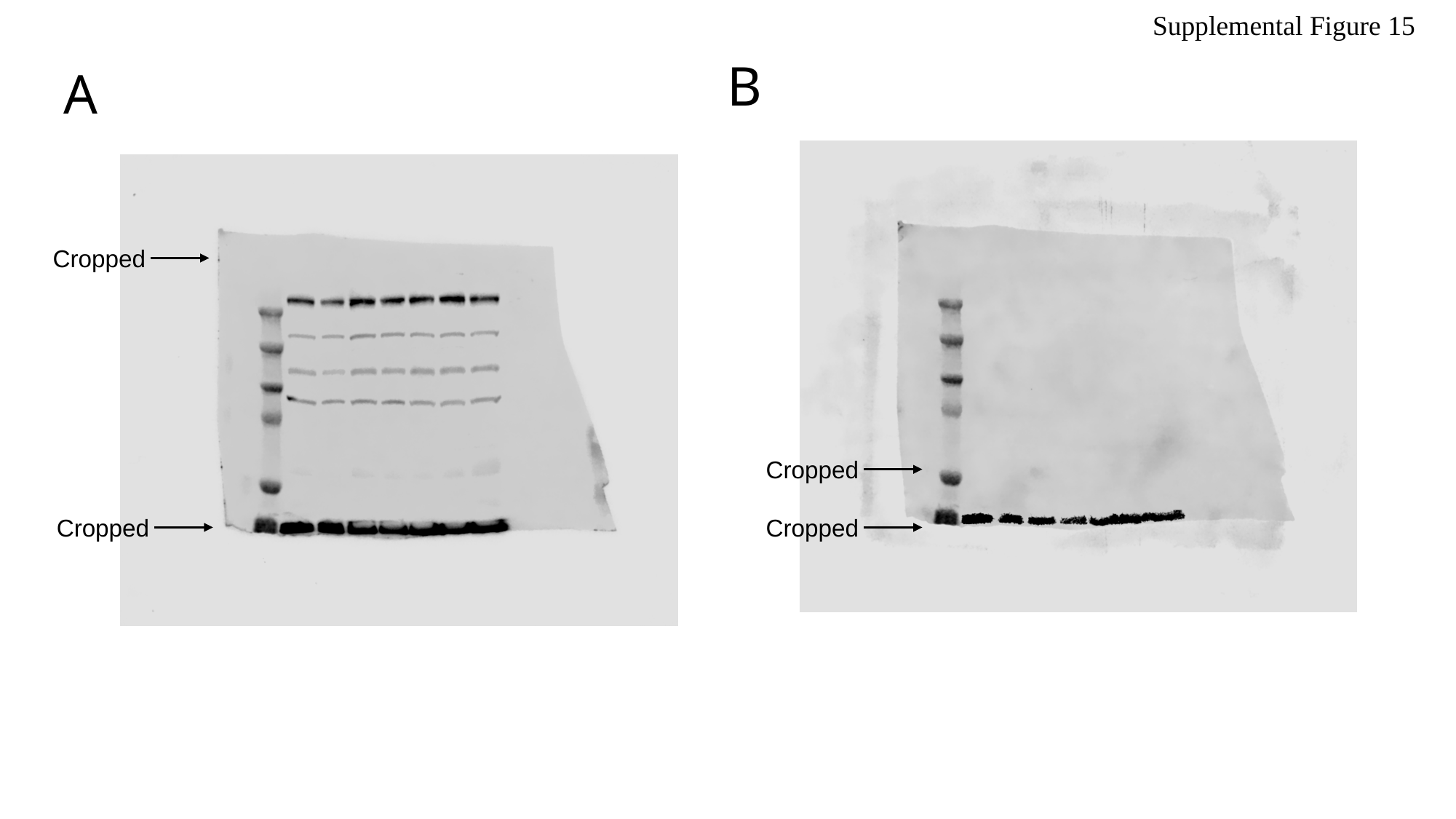

Supplemental Figure 15
B
A
 Cropped
 Cropped
 Cropped
 Cropped

### Slide 16
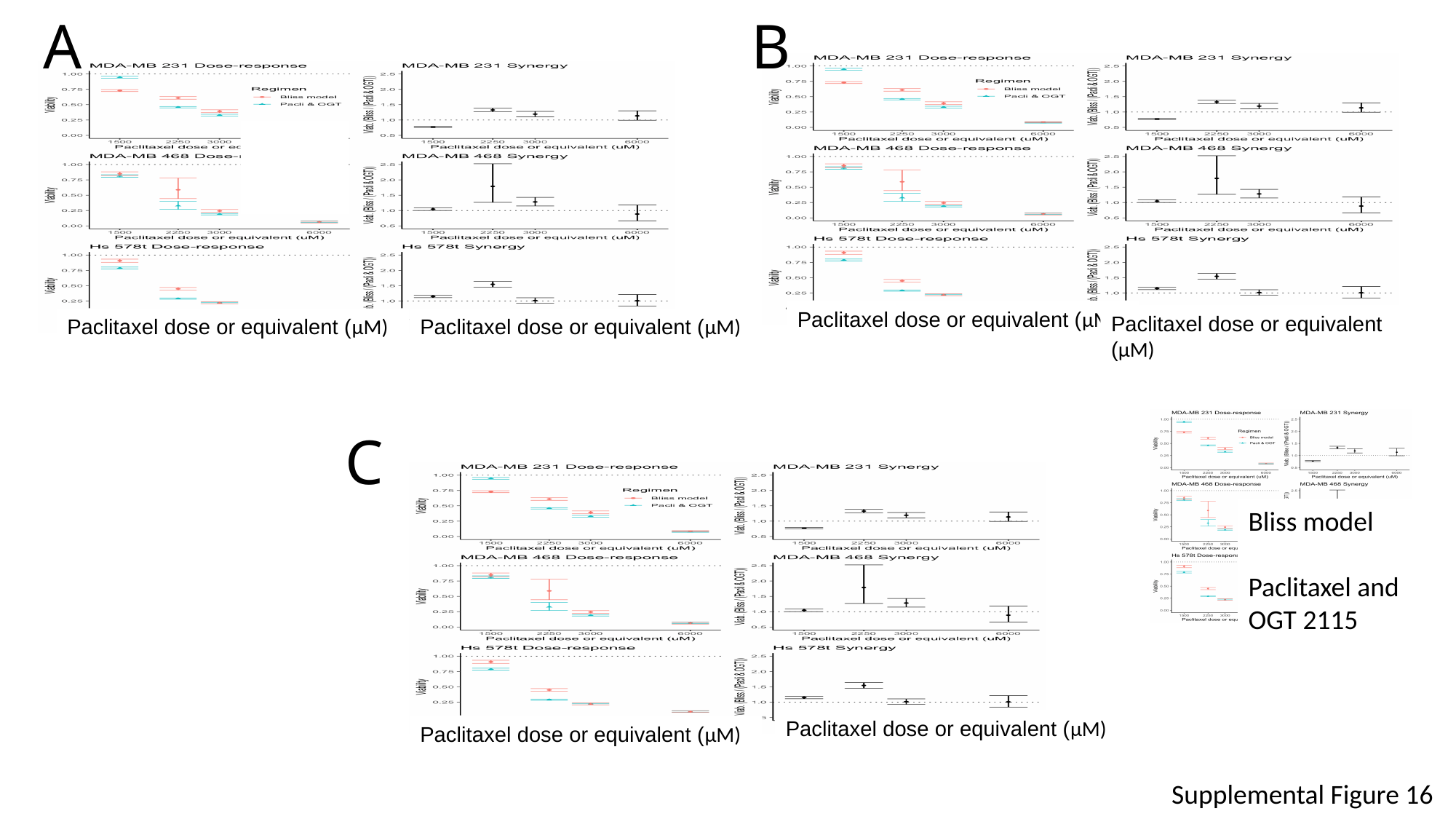

A
B
Paclitaxel dose or equivalent (µM)
Paclitaxel dose or equivalent (µM)
Paclitaxel dose or equivalent (µM)
Paclitaxel dose or equivalent (µM)
C
Bliss model
Paclitaxel and OGT 2115
Paclitaxel dose or equivalent (µM)
Paclitaxel dose or equivalent (µM)
Supplemental Figure 16

### Slide 17
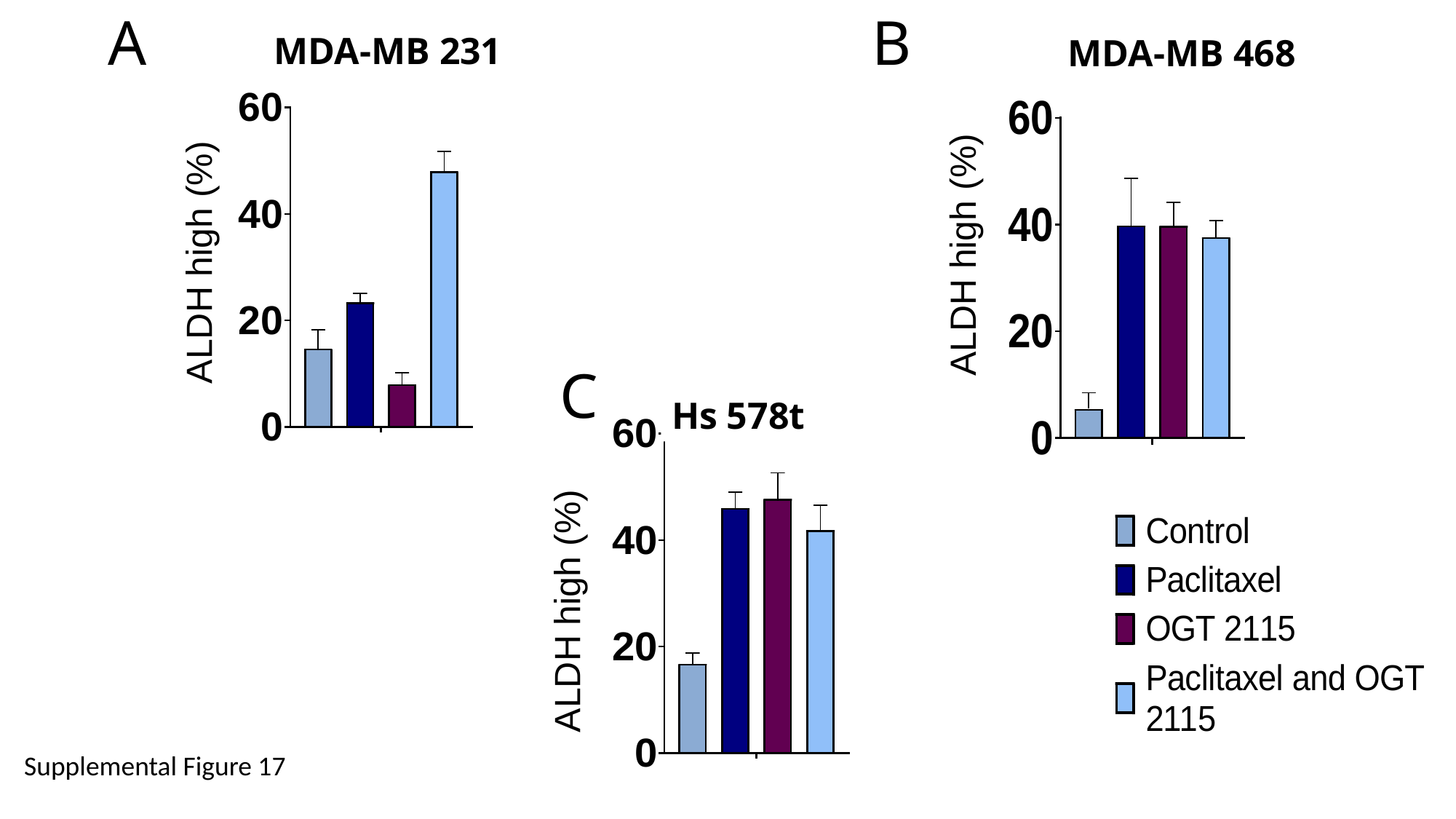

A
B
MDA-MB 231
MDA-MB 468
ALDH high (%)
ALDH high (%)
C
Hs 578t
ALDH high (%)
Supplemental Figure 17

### Slide 18
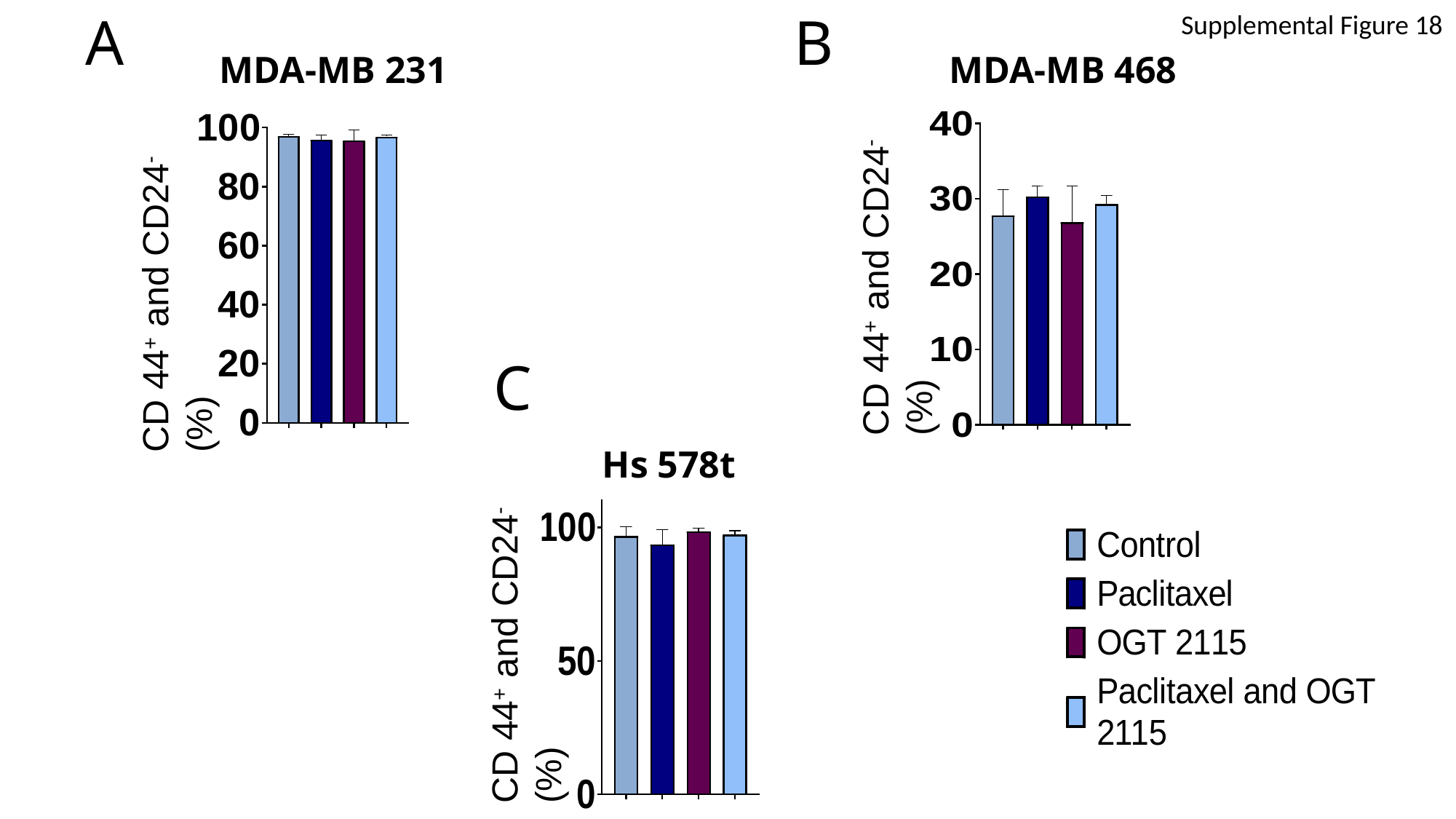

B
A
Supplemental Figure 18
MDA-MB 231
MDA-MB 468
CD 44^+^ and CD24^-^ (%)
CD 44^+^ and CD24^-^ (%)
C
CD 44^+^ and CD24^-^ (%)
Hs 578t

### Slide 19
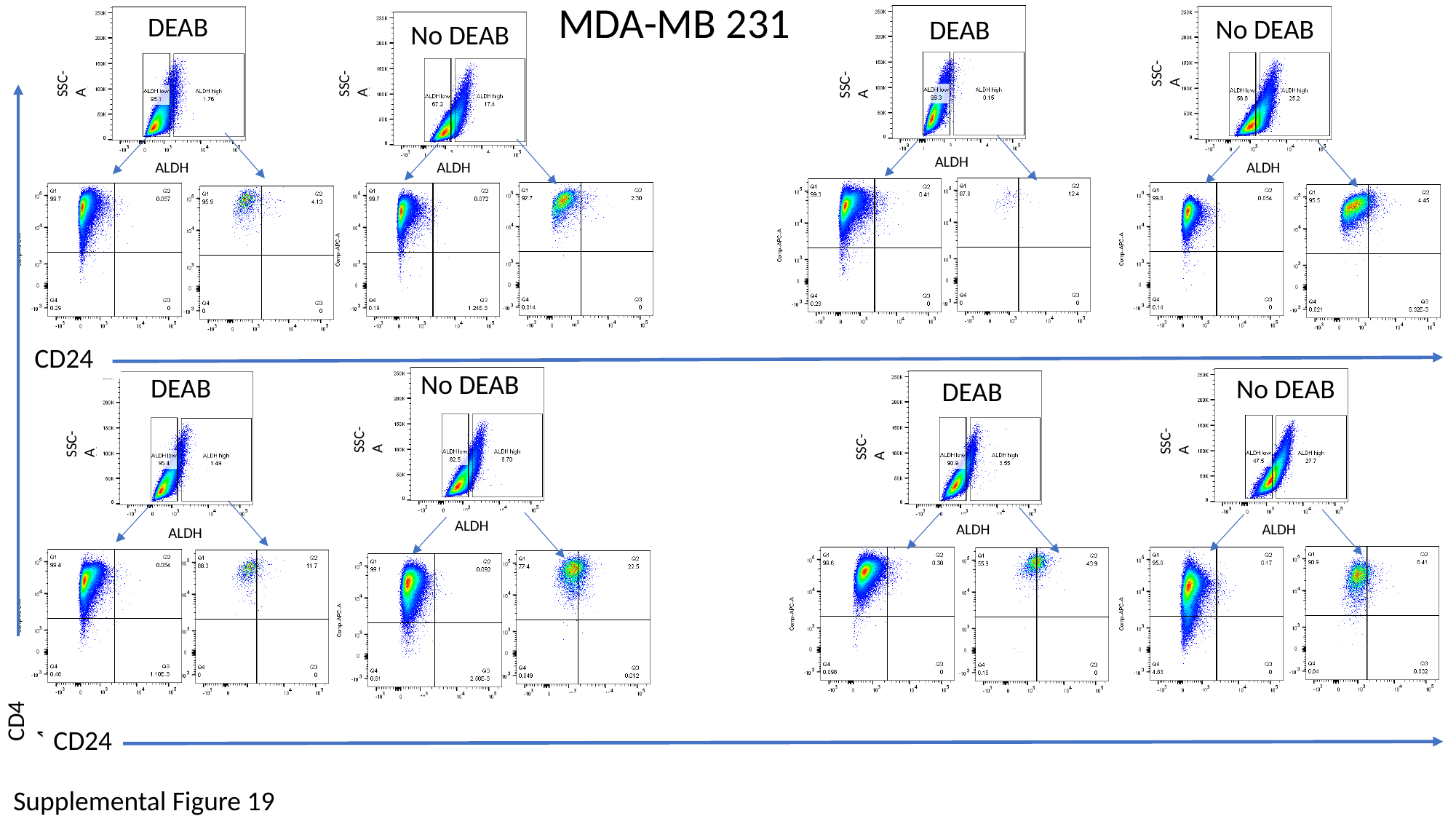

MDA-MB 231
DEAB
No DEAB
DEAB
No DEAB
SSC-A
SSC-A
SSC-A
SSC-A
SSC-A
SSC-A
ALDH
ALDH
ALDH
ALDH
CD24
No DEAB
DEAB
No DEAB
DEAB
SSC-A
SSC-A
SSC-A
SSC-A
SSC-A
SSC-A
ALDH
ALDH
ALDH
ALDH
CD44
CD24
Supplemental Figure 19

### Slide 20
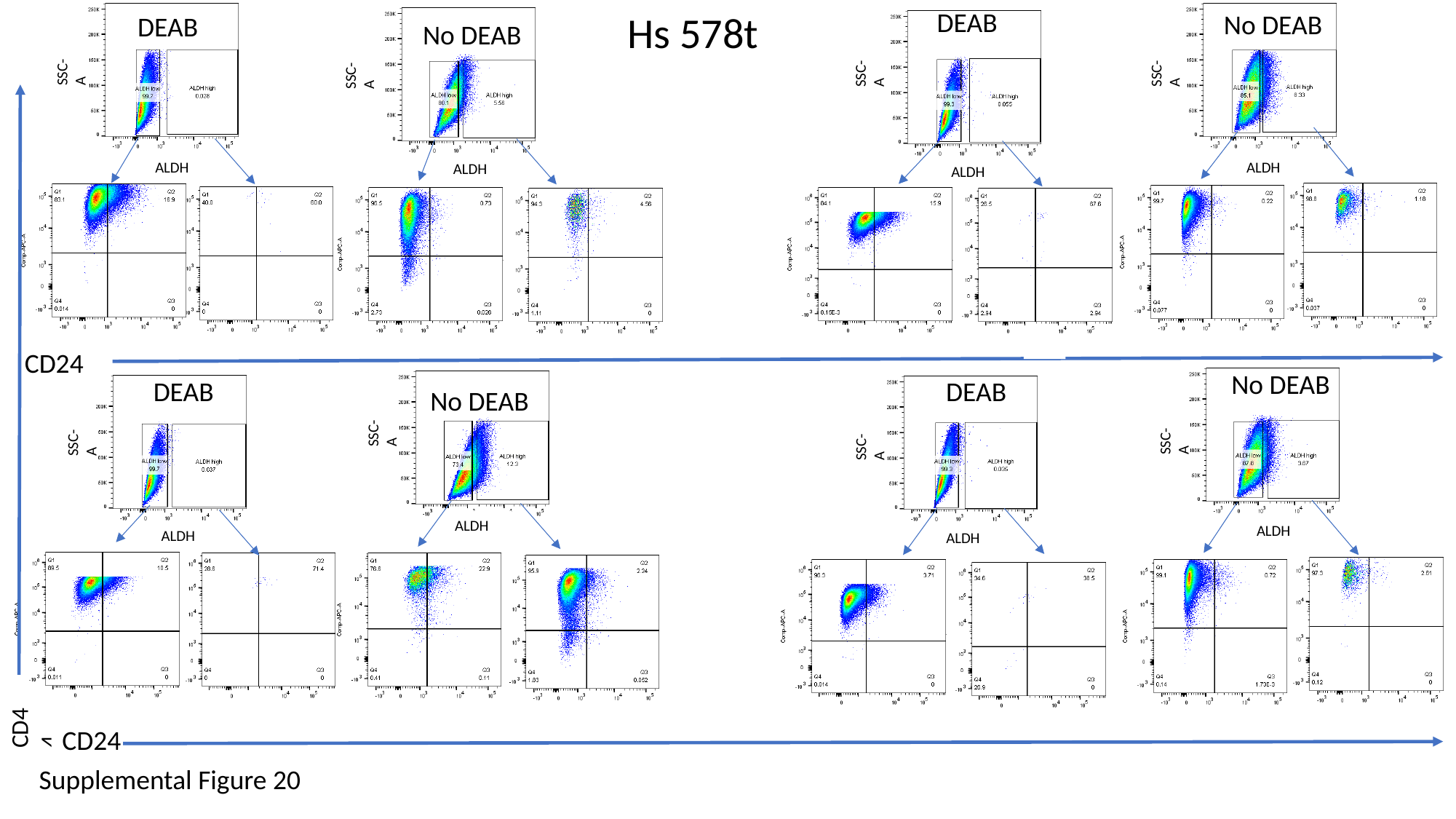

DEAB
Hs 578t
No DEAB
DEAB
No DEAB
SSC-A
SSC-A
SSC-A
SSC-A
ALDH
ALDH
ALDH
ALDH
CD24
No DEAB
DEAB
DEAB
No DEAB
SSC-A
SSC-A
SSC-A
SSC-A
ALDH
ALDH
ALDH
ALDH
CD44
CD24
Supplemental Figure 20

### Slide 21
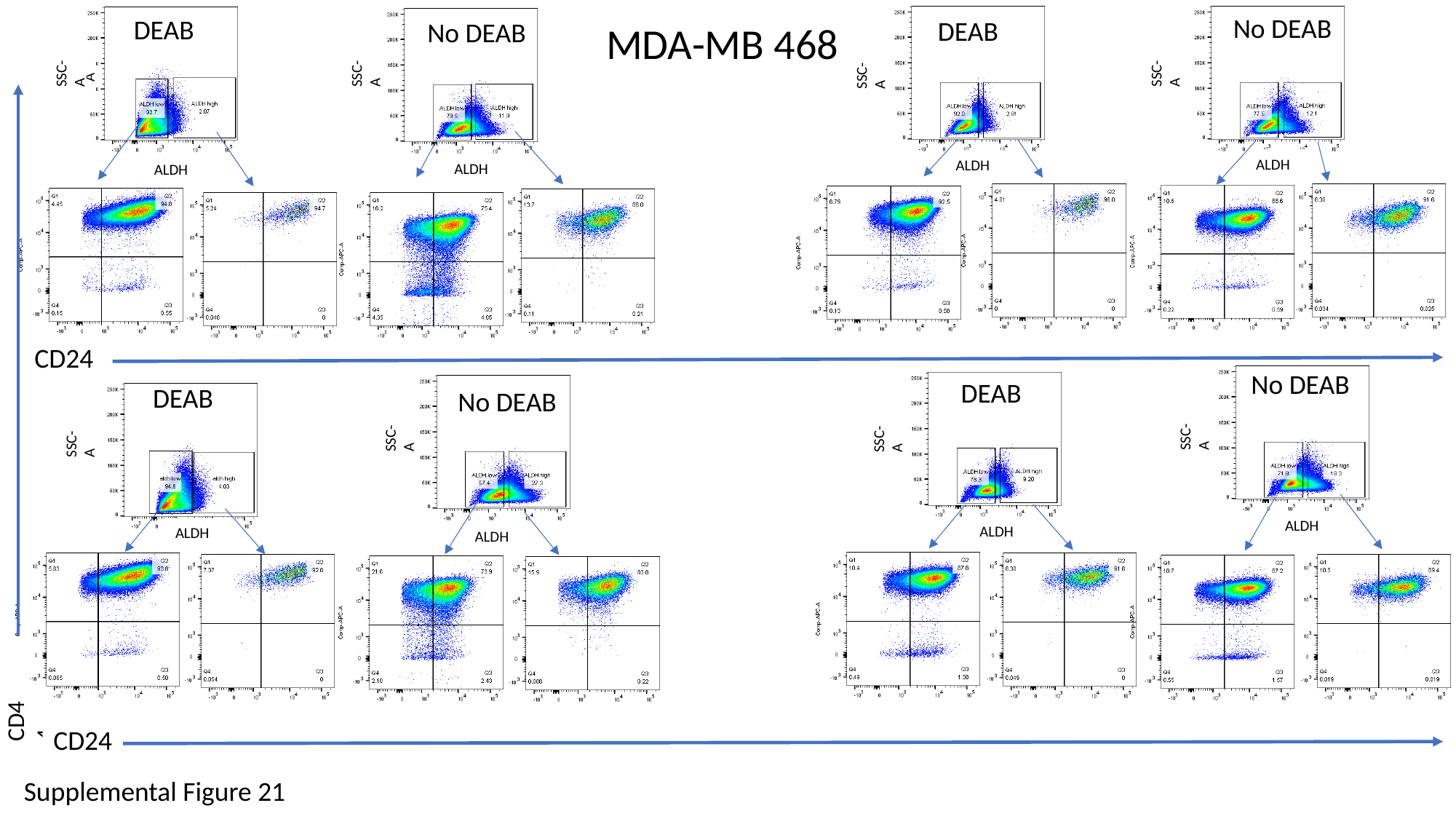

No DEAB
DEAB
DEAB
No DEAB
MDA-MB 468
SSC-A
SSC-A
SSC-A
SSC-A
SSC-A
ALDH
ALDH
ALDH
ALDH
CD24
No DEAB
DEAB
DEAB
No DEAB
SSC-A
SSC-A
SSC-A
SSC-A
ALDH
ALDH
ALDH
ALDH
CD44
CD24
Supplemental Figure 21

### Slide 22
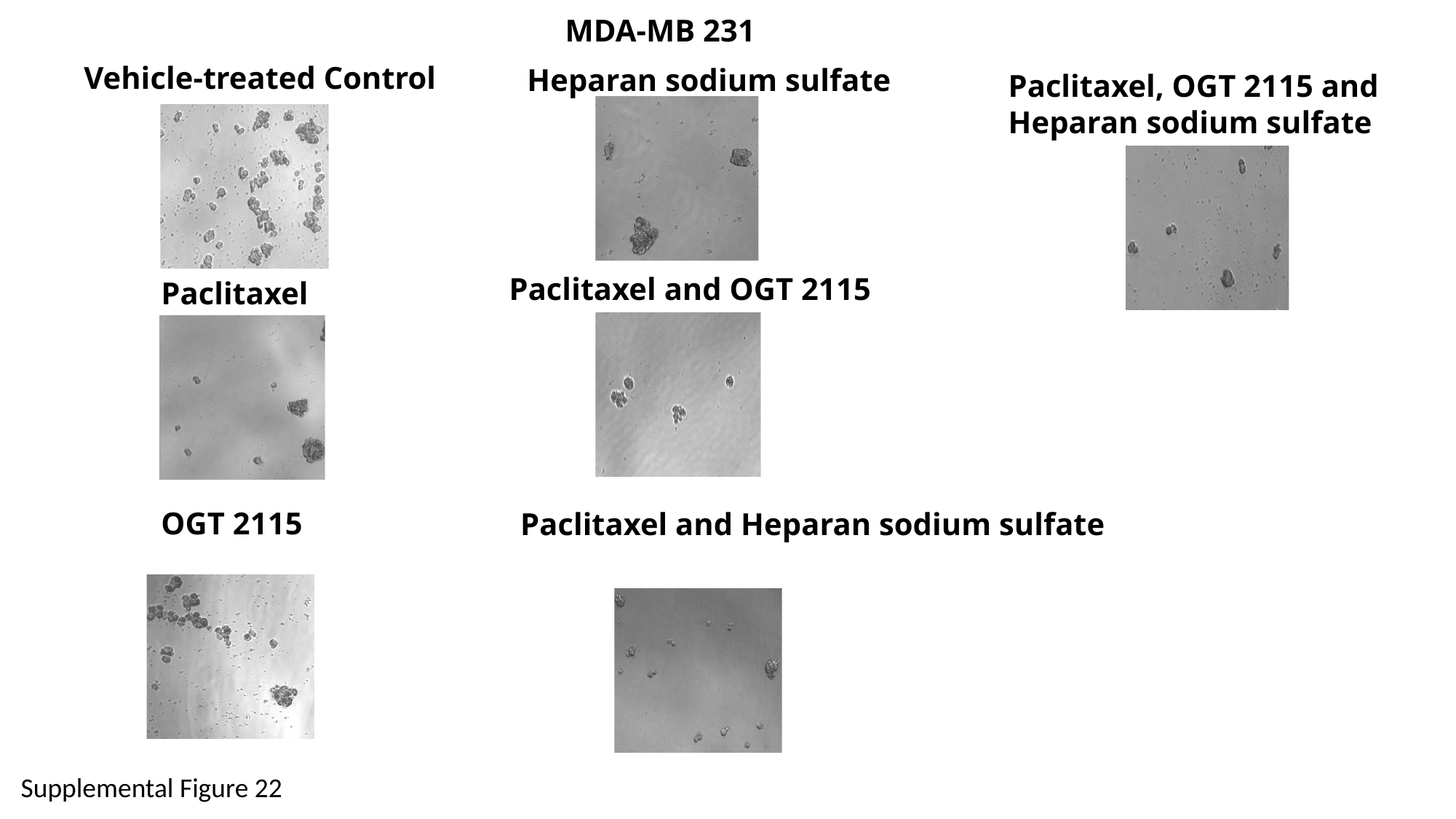

MDA-MB 231
Vehicle-treated Control
Heparan sodium sulfate
Paclitaxel, OGT 2115 and Heparan sodium sulfate
Paclitaxel and OGT 2115
Paclitaxel
OGT 2115
Paclitaxel and Heparan sodium sulfate
Supplemental Figure 22
Paclitaxel and Heparan sodium sulfate

### Slide 23
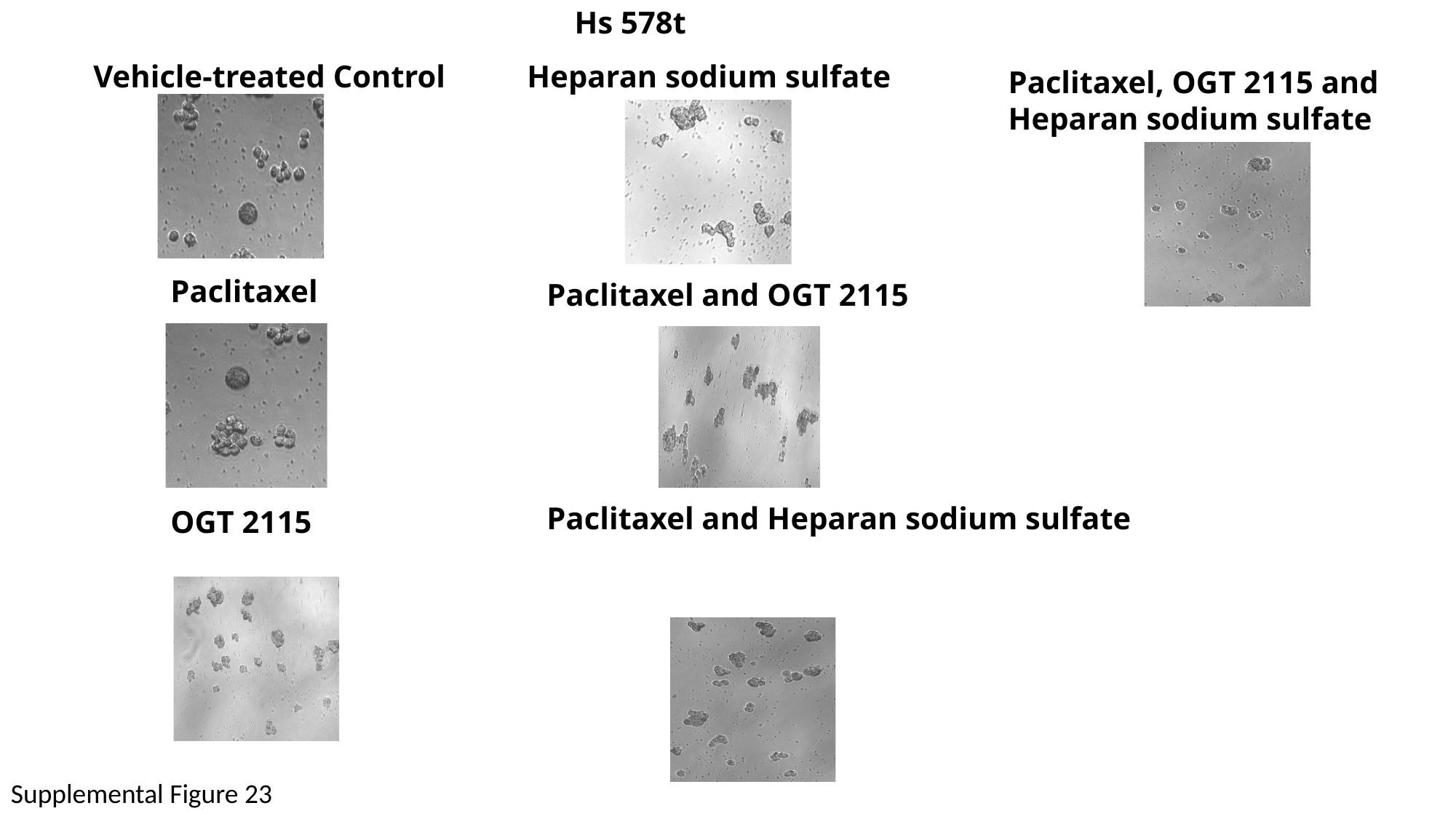

Hs 578t
Vehicle-treated Control
Heparan sodium sulfate
Paclitaxel, OGT 2115 and Heparan sodium sulfate
Paclitaxel
Paclitaxel and OGT 2115
Paclitaxel and Heparan sodium sulfate
OGT 2115
Supplemental Figure 23

### Slide 24
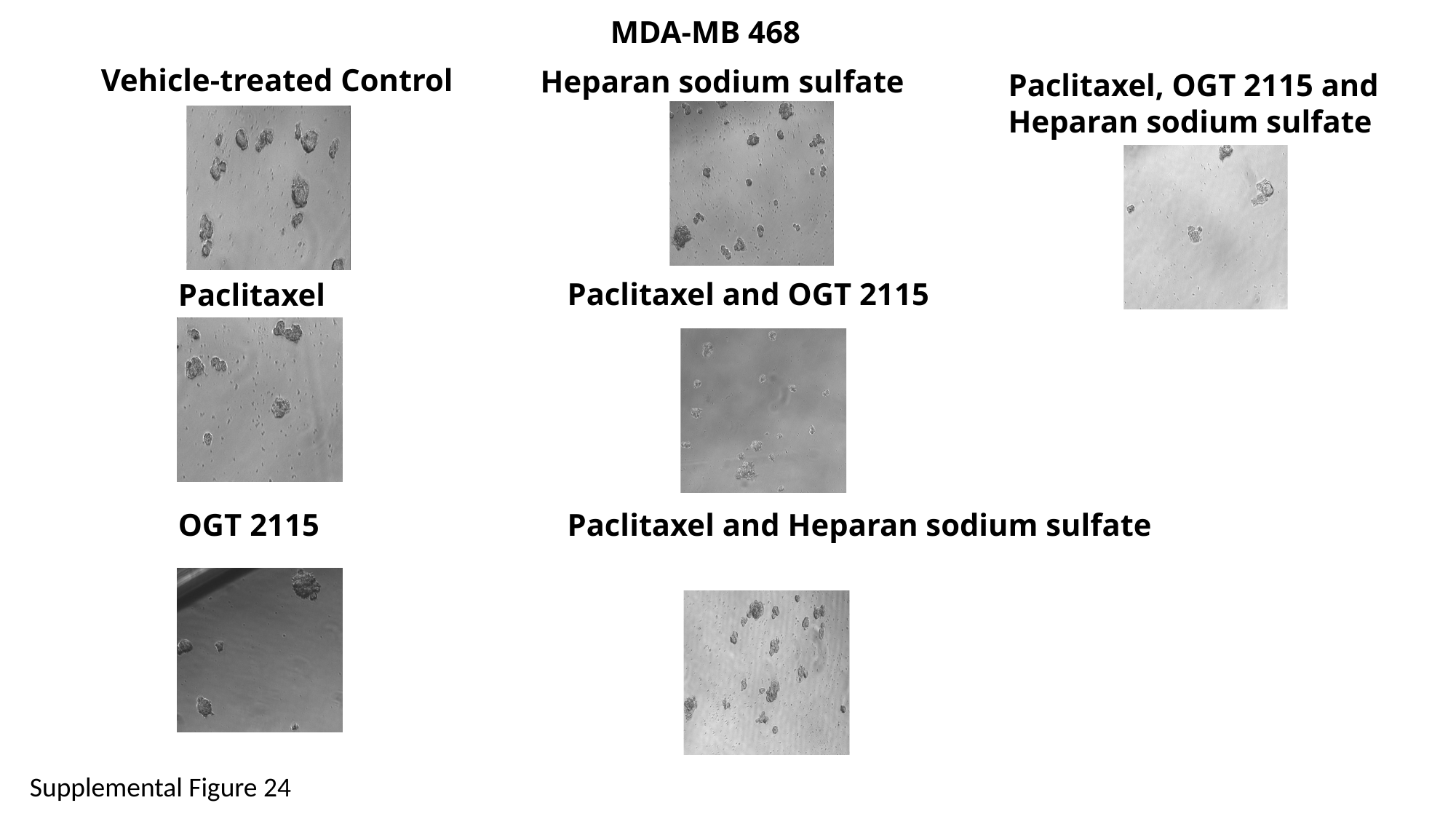

MDA-MB 468
Vehicle-treated Control
Heparan sodium sulfate
Paclitaxel, OGT 2115 and Heparan sodium sulfate
Paclitaxel and OGT 2115
Paclitaxel
OGT 2115
Paclitaxel and Heparan sodium sulfate
Supplemental Figure 24
