## Supplementary figures and images for "The role of heparan sulfate in enhancing the chemotherapeutic response in triple-negative breast cancer"

### Supplemental Table 1

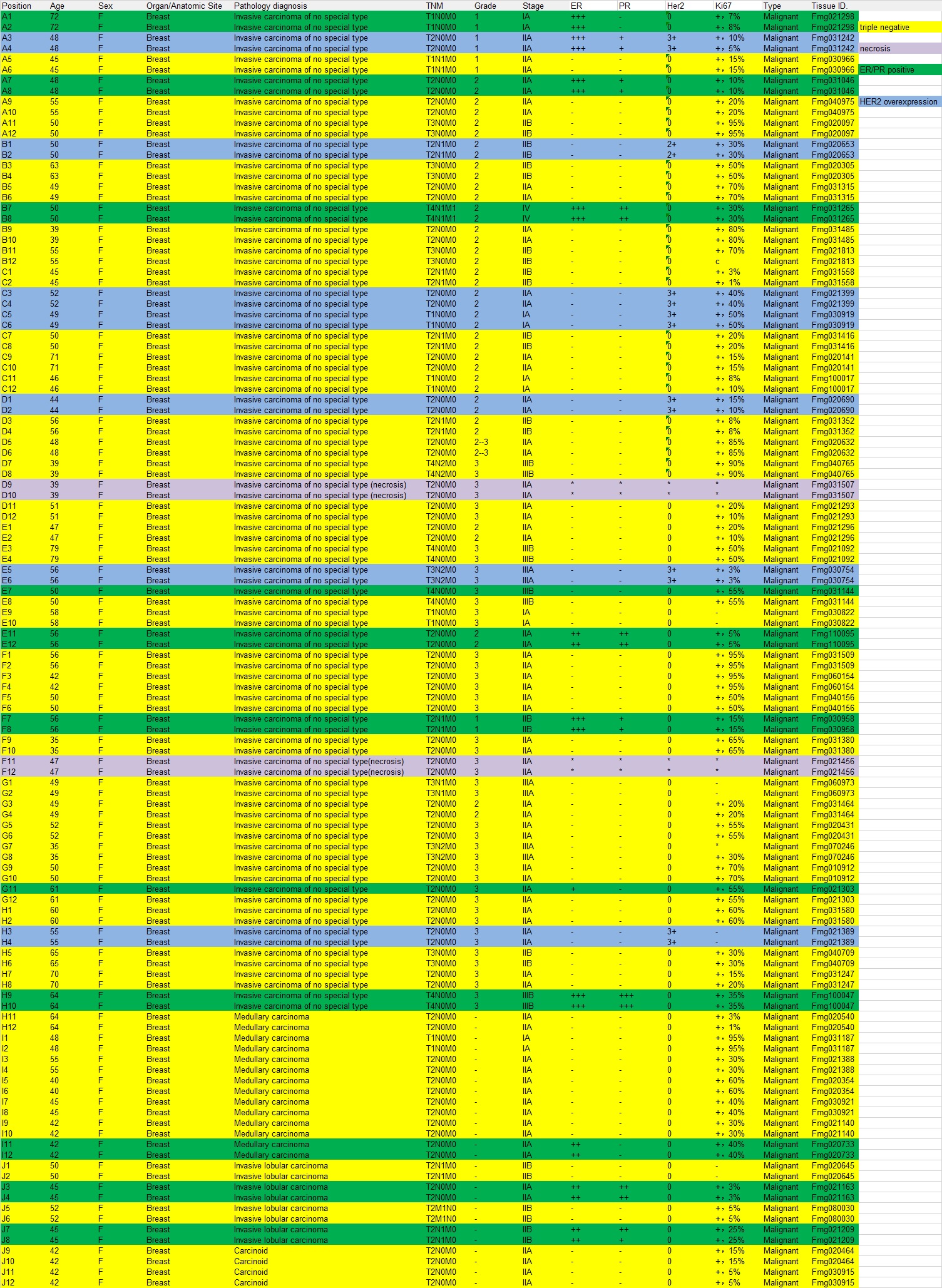
