## Supplemental Figure Legends for "The role of heparan sulfate in enhancing the chemotherapeutic response in triple-negative breast cancer"

**Supplemental Figure 1:** **Full western blot for heparanase from Figure 3A. (A)**The full, un-cropped western blot for heparanase from Figure 3A. **(B)** The full, un-cropped Ponceau S stain image for heparanase from Figure 3A.

**Supplemental Figure 2:** **Full western blot for heparanase from Figure 3B.** The full, un-cropped western blot for heparanase from Figure 3B.

**Supplemental Figure 3:** **Full western blot for heparanase from Figure 3C.** The full, un-cropped western blot for **(A)** heparanase and **(B)** GAPDH from Figure 3C.

**Supplemental Figure 4:** **Western blot analysis for TNBC and MCF-10A cells treated with paclitaxel or ATP**. TNBC MDA-MB 231, Hs 578t and MDA-MB 468 cells and nontumorigenic immortal mammary epithelial MCF-10A cells were treated with **(A)** paclitaxel (100 µM) for 6 hours or **(B)** ATP (500 µM) for 48 hours and 5 μM cell supernatants were probed heparanase. Ponceau S staining was performed immediately after transfer onto the nitrocellulose membranes to demonstrate loading of the proteins. Similar results were obtained in biological replicate experiment.

**Figure 6: Heparanase and Syndecan-1. (A)** Equal amounts of untreated and treated MDA-MB 468 cell lysates (100 µg) were probed for syndecan-1. Lane order for the blot is molecular weight marker, untreated cell lysate, lysates treated with: 50 units of heparanase, 20 µM OGT 2115, 20 µM OGT 2115 and 50 units of heparanase, 40 µM OGT 2115, 40 µM OGT 2115 and 50 units of heparanase and 100 units of heparin. (B) The densitometric analyses of the bands were calculated. The student’s t-test was performed to determine significance. * represents p<0.05 and ** represents p<0.01 when comparing expression in control MDA-MB 468 cell lysates to the expression in treated MDA-MB 468 cell lysates.

**Supplemental Figure 5:** **Full western blot for heparanase from Figure 4A. (A)** The full, un-cropped western blot for heparanase from Supplemental Figure 4A. **(B)** The full, un-cropped Ponceau S stain image for heparanase from Supplemental Figure 4A.

**Supplemental Figure 6:** **Full western blot for heparanase from Figure 4B. (A)** The full, un-cropped western blot for heparanase from Supplemental Figure 4B. **(B)** The full, un-cropped Ponceau S stain image for heparanase from Supplemental Figure 4B.

**Supplemental Figure 7:** **Heparan sulfate protein expression analysis.** Histogram obtained from flow cytometry analysis of heparan sulfate on the cell surface of the TNBC MDA-MB 231, Hs 578t and MDA-MB 468 cell lines and non-tumorigenic immortal epithelial mammary MCF-10A cells. **(B)** Basal heparan sulfate expression was examined in the supernatants of TNBC cells and MCF-10A via ELISA analysis with MDA-MB 468 expressing the most. Standard deviation was calculated from three independent experiments performed in triplicate. The student’s t-test was performed to determine significance with * represents p<0.05 and ** represents p<0.01 the protein expression in MCF-10A to the protein expressions in the TNBC cell lines.

**Supplemental Figure 8: Immunohistochemistry for heparan sulfate staining. (A)** Normal breast tissue on slides (2) stained for heparan sulfate. **(B)** Ductal carcinoma in situ (DCIS) tissue on slides (3) stained for heparan sulfate. **(C)** Slide key for the breast cancer tissue array orientation.

**Supplemental Figure 9: Immunohistochemistry for heparanase staining. (A)** Normal breast tissue on slides (2) stained for heparanase. **(B)** Ductal carcinoma in situ (DCIS) tissue on slides (3) stained for heparanase.

**Supplemental Figure 10:** **Statistical analysis for heparan sulfate immunohistochemistry comparing different grades of breast cancer to normal breast tissue: (A)** Pairwise comparisons using Dunn’s test showed that there was a significant difference between breast cancer Grade 1 and normal (p = 0.0030), Grade 1 and DCIS (p = 0.0203), Grade 2 and normal (p = 0.0434), Grade 3 and normal (p = 0.0019), and Grade 3 and DCIS (p = 0.0222), in the average % of heparan sulfate positively stained cells. No other differences were significantly different. **(B)** Pairwise comparisons using Dunn’s test indicated that there was a significant difference between breast cancer Grade 1 and normal (p = 0.0031), Grade 1 and DCIS (p = 0.0207), Grade 2 and normal (p = 0.0442), Grade 3 and normal (p = 0.0018), and Grade 3 and DCIS (p = 0.0217), in the average % of heparan sulfate weakly stained cells. No other differences were significantly different. **(C)** Pairwise comparisons using Dunn’s test indicated that there was a significant difference between breast cancer Grade 1 and normal (p = 0.0006), Grade 1 and DCIS (p = 0.0064), Grade 2 and normal (p = 0.008), Grade 2 and DCIS (p = 0.0141), Grade 3 and normal (p = 0.0024), and Grade 3 and DCIS (p = 0.0315), in the average % of heparan sulfate moderately stained cells. No other differences were significantly different. **(D)** Pairwise comparisons using Dunn’s test indicated that there was a significant difference between breast cancer Grade 1 and normal (p = 0.0007), Grade 1 and DCIS (p = 0.0151), Grade 2 and normal (p = 0.0001), Grade 2 and DCIS (p = 0.0044), Grade 3 and normal (p = 0.0001), and Grade 3 and DCIS (p = 0.0028), in the average % of heparan sulfate strongly stained cells. No other differences were significantly different. **(E)** Pairwise comparisons using Dunn’s test demonstrated that there was a significant difference between breast cancer Grade 1 and normal (p = 0.0030), Grade 1 and DCIS (p = 0.0203), Grade 2 and normal (p = 0.0434), Grade 3 and normal (p = 0.0019), and Grade 3 and DCIS (p = 0.0222), in the average % of negative heparan sulfate stained cells. No other differences were significantly different.

**Supplemental Figure 11:** **Statistical analysis for heparan sulfate immunohistochemistry comparing % Ki 67 expression levels.** Kruskal-Wallis test demonstrated that there was no significant difference: **(A)** in the average % of heparan sulfate positively stained cells amongst different % Ki67 expression levels, **(B)** in the average % of heparan sulfate weakly stained cells amongst different % Ki67 expression levels, **(C)** in the average % of heparan sulfate moderately stained cells amongst different % Ki67 expression levels, **(D)** in the average % of heparan sulfate strongly stained cells amongst different % Ki67 expression levels and **(E)** in the average % of negative heparan sulfate stained cells amongst different % Ki67 expression levels.

**Supplemental Figure 12:** **Statistical analysis for heparanase immunohistochemistry comparing different grades of breast cancer and normal breast tissue.** Kruskal-Wallis test showed that there was no significant difference: **(A)** in the average % of heparanase positively stained cells amongst different cancer grades and normal breast tissue, **(B)** in the average % of heparanase weakly stained cells amongst different cancer grades and normal breast tissue, **(C)** in the average % of heparanase moderately stained cells in the tumor amongst different cancer grades and normal breast tissue, **(D)** in the average % of heparanase strongly stained cells in the tumor amongst different cancer grades and normal breast tissue and **(E)** in the average % of negative heparanase stained cells in the tumor amongst different cancer grades and normal breast tissue.

**Supplemental Figure 13:** **Statistical analysis for heparanase immunohistochemistry comparing % Ki67 expression levels.** Kruskal-Wallis test demonstrated that there was no significant difference: **(A)** in the average % of heparanase positively stained cells in the tumor amongst different % Ki67 expression levels, **(B)** in the average % of heparanase weakly stained cells in the tumor amongst different % Ki67 expression levels, **(C)** in the average % of heparanase moderately stained cells in the tumor amongst different % Ki67 expression levels, **(D)** in the average % of heparanase moderately stained cells in the tumor amongst different % Ki67 expression levels and **(E)** in the average % of negative heparanase stained cells in the tumor amongst different % Ki67 expression levels.

**Figure 14: Heparanase and Syndecan-1. (A**) Equal amounts of untreated and treated MDA-MB 468 cell lysates (100 µg) were probed for syndecan-1. Lane order for the blot is molecular weight marker, untreated cell lysate, lysates treated with: 50 units of heparanase, 20 µM OGT 2115, 20 µM OGT 2115 and 50 units of heparanase, 40 µM OGT 2115, 40 µM OGT 2115 and 50 units of heparanase and 100 units of heparin. **(B)** The densitometric analyses of the bands were calculated. The student’s t-test was performed to determine significance. * represents p<0.05 and ** represents p<0.01 when comparing expression in control MDA-MB 468 cell lysates to the expression in treated MDA-MB 468 cell lysates.

**Supplemental Figure 15:** **Full western blot for syndecan-1 expression from Figure 6. (A)** The full, un-cropped western blot for syndecan-1 from Supplemental Figure 14. **(B)** The full, un-cropped western blot for GAPDH for syndecan-1 from Supplemental Figure 14.

**Supplemental Figure 16:** **Statistical analysis of treated cells with sulfatase inhibitor OGT 2115 and chemotherapeutic agent paclitaxel.** Dose response and synergy graphs are displayed for TNBC **(A)** MDA-MB 231 **(B)** Hs 578t and **(C)** MDA-MB 468 with increasing concentrations of paclitaxel, OGT 2115 or the co-treatment administered for 48 hours. For the dose response graphs, the bliss model (orange) and the co-treatment (paclitaxel and OGT 2115) are shown. There is some synergy (<0.1-1.0) for one dose combination for MDA-MB 231 cells while there were some drug dose combinations that were additive (1-1.2) for Hs 578t and MDA-MB 468 cells. These graphs were obtained from three independent experiments performed in triplicate.

**Supplemental Figure 17:** **ALDH expression in TNBC cells.** The % of cells is shown for **(A)** MDA-MB 231 **(B)** MDA-MB 468 and **(C)** Hs 578t that are considered ALDH high. TNBCs treated with paclitaxel alone produced the most cells that expressed ALDH at high concentrations. Standard deviation was calculated from three independent experiments performed in triplicate. 1-way ANOVA with Tukey’s HSD was applied to ascertain significance. * represents p<0.05 and ** represents p<0.01 when comparing paclitaxel to paclitaxel and OGT 2115.

**Supplemental Figure 18: CD44 and CD24 expressions in TNBC cells.** The % of cells is shown for **(A)** MDA-MB 231 **(B)** MDA-MB 468 and **(C)** Hs 578t that are considered CD44 positive and CD24 negative. MDA-MB 231 and Hs 578t cells highly express CD44 under all drug conditions; whereas, MDA-MB 468 cells treated with vehicle or paclitaxel expressed more CD44 than those treated with OGT 2115 or the co-treatment of OGT 2115 and paclitaxel. Standard deviation was calculated from three independent experiments performed in triplicate. 1-way ANOVA with Tukey’s HSD was applied to ascertain significance. * represents p<0.05 and ** represents p<0.01 when comparing paclitaxel to paclitaxel and OGT 2115.

**Supplemental Figure 19:** **Cancer-initiating cell dot plots for OGT 2115 and paclitaxel-treated MDA-MB 231 cells**. Dot plots examining the cancer-initiating cells that are ALDH high, CD44 positive, and CD24 negative are presented for treated MDA-MB 231 cells with paclitaxel, sulfatase inhibitor OGT 2115, or both drug agents. Diethylaminobenzaldehyde (DEAB) is a specific inhibitor of ALDH, that can be used as a background fluorescence control. Three independent experiments were performed in triplicate.

**Supplemental Figure 20:** **Cancer-initiating cell dot plots for OGT 2115 and paclitaxel-treated Hs 578t cells**. Dot plots examining the cancer-initiating cells that are ALDH high, CD44 positive, and CD24 negative are presented for treated Hs 578t cells with paclitaxel, sulfatase inhibitor OGT 2115, or both drug agents. DEAB is used as a background fluorescence control. Three independent experiments were performed in triplicate.

**Supplemental Figure 21:** **Cancer-initiating cell dot plots for OGT 2115 and paclitaxel-treated MDA-MB 468**. Dot plots examining the cancer-initiating cells that are ALDH high, CD44 positive, and CD24 negative are presented for treated MDA-MB 468 cells with paclitaxel, sulfatase inhibitor OGT 2115, or both drug agents. DEAB is used as a background fluorescence control. Three independent experiments were performed in triplicate.

**Supplemental Figure 22:** **Tumorsphere efficiency assay images for treated MDA-MB 231 cells**. Tumorsphere images obtained from the Etaluma™ Lumascope 620 (10X) are displayed for each treatment of MDA-MB 231 cells with paclitaxel, OGT 2115, heparan sodium sulfate, or the different combinations. Three independent experiments were performed in triplicate.

**Supplemental Figure 23:** **Tumorsphere efficiency assay images for treated MDA-MB 468 cells**. Tumorsphere images obtained from the Etaluma™ Lumascope 620 (10X) are displayed for each treatment of MDA-MB 468 cells with paclitaxel, OGT 2115, heparan sodium sulfate, or the different combinations. Three independent experiments were performed in triplicate.

**Supplemental Figure 24:** **Tumorsphere efficiency assay images for treated Hs 578t cells**. Tumorsphere images obtained from the Etaluma™ Lumascope 620 (10X) are displayed for each treatment of Hs 578t cells with paclitaxel, OGT 2115, heparan sodium sulfate, or the different combinations. Three independent experiments were performed in triplicate.
